## Supplementary material for "Does diversity beget diversity in microbiomes?": File S1

### Full GLMMs output for EMP data

Significant models (likelihood-ratio test, p<0.05)

Naïma Madi

2019-09-15

#### 1. DBD across different taxonomic ratios

##### 1.1. ASV:Genus

```
## Generalized linear mixed model fit by maximum likelihood (Laplace
##   Approximation) [glmerMod]
##  Family: poisson  ( log )
## Formula: nb_ASV ~ nb_genus + (nb_genus | genus_type/empo_3) + (nb_genus |
##     empo_3:PI) + (nb_genus - 1 | sample)
##    Data: datsc1
## Control: glmerControl(optimizer = "bobyqa", optCtrl = list(maxfun = 1e+05))
##
##       AIC       BIC    logLik  deviance  df.resid
##  290657.5  290771.1 -145316.7  290633.5     95593
##
## Scaled residuals:
##     Min      1Q  Median      3Q     Max
## -3.7143 -0.3896 -0.1390  0.2107 12.0218
##
## Random effects:
##  Groups            Name        Variance Std.Dev. Corr
##  empo_3:genus_type (Intercept) 0.056526 0.23775
##                    nb_genus    0.020192 0.14210  0.04
##  sample            nb_genus    0.004483 0.06696
##  genus_type        (Intercept) 0.069547 0.26372
##                    nb_genus    0.005417 0.07360  0.56
##  empo_3:PI         (Intercept) 0.016527 0.12856
##                    nb_genus    0.013038 0.11418  0.06
## Number of obs: 95605, groups:
## empo_3:genus_type, 7857; sample, 1995; genus_type, 1128; empo_3:PI, 117
##
## Fixed effects:
##             Estimate Std. Error z value Pr(>|z|)
## (Intercept)  0.31772    0.01822  17.441  < 2e-16 ***
## nb_genus     0.09075    0.01567   5.792 6.95e-09 ***
## ---
## Signif. codes:  0 '***' 0.001 '**' 0.01 '*' 0.05 '.' 0.1 ' ' 1
```

nb_ASV=ASV:genus, nb_genus=number of non-focal genera, genus_type=Lineage, empo_3=Environment, PI=laboratory that submitted the data, sample=EMP sample ID.

##### 1.2. Genus:Family

```
## Generalized linear mixed model fit by maximum likelihood (Laplace
##   Approximation) [glmerMod]
##  Family: poisson  ( log )
## Formula:
## nb_genus ~ nb_family + (nb_family | family_type/empo_3) + (nb_family |
##     empo_3:PI)
##    Data: datsc
## Control: glmerControl(optimizer = "bobyqa", optCtrl = list(maxfun = 1e+05))
##
##       AIC       BIC    logLik  deviance  df.resid
##  277313.9  277419.8 -138646.0  277291.9    111922
##
## Scaled residuals:
##     Min      1Q  Median      3Q     Max
## -2.0954 -0.2301 -0.0484  0.0452  6.2366
##
## Random effects:
##  Groups             Name        Variance Std.Dev. Corr
##  empo_3:family_type (Intercept) 0.022386 0.14962
##                     nb_family   0.005039 0.07099  0.08
##  family_type        (Intercept) 0.049366 0.22218
##                     nb_family   0.005063 0.07115  0.74
##  empo_3:PI          (Intercept) 0.001811 0.04256
##                     nb_family   0.001532 0.03914  -0.38
## Number of obs: 111933, groups:
## empo_3:family_type, 4832; family_type, 458; empo_3:PI, 117
##
## Fixed effects:
##             Estimate Std. Error z value Pr(>|z|)
## (Intercept) 0.167032   0.013531  12.344  < 2e-16 ***
## nb_family   0.047326   0.008007   5.911 3.41e-09 ***
## ---
## Signif. codes:  0 '***' 0.001 '**' 0.01 '*' 0.05 '.' 0.1 ' ' 1
```

nb_genus=genus:family, nb_family=number of non-focal families, family_type=Lineage, empo_3=Environment, PI=laboratory that submitted the data.

##### 1.3. Family:Order

```
## Generalized linear mixed model fit by maximum likelihood (Laplace
##   Approximation) [glmerMod]
##  Family: poisson  ( log )
## Formula: nb_family ~ nb_order + (nb_order | order_type/empo_3) + (nb_order -
##     1 | empo_3) + (nb_order | empo_3:PI)
##    Data: datsc
## Control: glmerControl(optimizer = "bobyqa", optCtrl = list(maxfun = 1e+05))
##
##       AIC       BIC    logLik  deviance  df.resid
```

```
##  261737.7  261852.4 -130856.8  261713.7    105182
##
## Scaled residuals:
##     Min      1Q  Median      3Q     Max
## -2.6704 -0.1386 -0.0131  0.0567  6.5306
##
## Random effects:
##  Groups            Name        Variance  Std.Dev. Corr
##  empo_3:order_type (Intercept) 0.0210015 0.14492
##                    nb_order    0.0084763 0.09207  -0.07
##  order_type        (Intercept) 0.0837311 0.28936
##                    nb_order    0.0087631 0.09361  0.74
##  empo_3:PI         (Intercept) 0.0078208 0.08844
##                    nb_order    0.0112401 0.10602  0.69
##  empo_3            nb_order    0.0005462 0.02337
## Number of obs: 105194, groups:
## empo_3:order_type, 3781; order_type, 363; empo_3:PI, 117; empo_3, 17
##
## Fixed effects:
##             Estimate Std. Error z value Pr(>|z|)
## (Intercept)  0.19068    0.02149   8.875  < 2e-16 ***
## nb_order     0.11918    0.01702   7.001 2.54e-12 ***
## ---
## Signif. codes:  0 '***' 0.001 '**' 0.01 '*' 0.05 '.' 0.1 ' ' 1
```

nb_family=family:order, nb_order=number of non-focal orders, order_type=Lineage, empo_3=Environment, PI=laboratory that submitted the data.

##### 1.4. Order:Class

```
## Generalized linear mixed model fit by maximum likelihood (Laplace
##   Approximation) [glmerMod]
##  Family: poisson  ( log )
## Formula:
## nb_order ~ nb_classe + (nb_classe | classe_type/empo_3) + (nb_classe -
##     1 | empo_3) + (nb_classe | empo_3:PI)
##    Data: datsc
## Control: glmerControl(optimizer = "bobyqa", optCtrl = list(maxfun = 1e+05))
##
##      AIC      BIC   logLik deviance df.resid
## 189330.9 189441.7 -94653.5 189306.9    75369
##
## Scaled residuals:
##     Min      1Q  Median      3Q     Max
## -2.2595 -0.1457 -0.0179  0.0429  4.1586
##
## Random effects:
##  Groups             Name        Variance  Std.Dev. Corr
##  empo_3:classe_type (Intercept) 0.0134159 0.11583
##                     nb_classe   0.0061698 0.07855  -0.01
##  classe_type        (Intercept) 0.1122675 0.33506
##                     nb_classe   0.0199721 0.14132  0.70
##  empo_3:PI          (Intercept) 0.0008366 0.02892
##                     nb_classe   0.0026542 0.05152  -0.29
```

```
##  empo_3             nb_classe   0.0025565 0.05056
## Number of obs: 75381, groups:
## empo_3:classe_type, 2383; classe_type, 230; empo_3:PI, 117; empo_3, 17
##
## Fixed effects:
##             Estimate Std. Error z value Pr(>|z|)
## (Intercept)  0.17490    0.02499   6.998 2.61e-12 ***
## nb_classe    0.10872    0.01996   5.447 5.13e-08 ***
## ---
## Signif. codes:  0 '***' 0.001 '**' 0.01 '*' 0.05 '.' 0.1 ' ' 1
```

nb_order=order:class, nb_classe=number of non-focal classes, classe_type=Lineage, empo_3=Environment, PI=laboratory that submitted the data.

##### 1.5. Class:Phylum

```
## Loading required package: lme4

## Loading required package: Matrix

## Generalized linear mixed model fit by maximum likelihood (Laplace
##   Approximation) [glmerMod]
##  Family: poisson  ( log )
## Formula:
## nb_class ~ nb_phylum + (nb_phylum | phylum_type/empo_3) + (nb_phylum |
##     empo_3:PI) + (nb_phylum - 1 | empo_3)
##    Data: datsc
## Control: glmerControl(optimizer = "bobyqa")
##
##      AIC      BIC   logLik deviance df.resid
##  99420.6  99521.7 -49698.3  99396.6    33702
##
## Scaled residuals:
##     Min      1Q  Median      3Q     Max
## -2.6993 -0.2934 -0.0350  0.2070  4.7465
##
## Random effects:
##  Groups             Name        Variance Std.Dev. Corr
##  empo_3:phylum_type (Intercept) 0.026141 0.16168
##                     nb_phylum   0.014168 0.11903  -0.36
##  empo_3:PI          (Intercept) 0.009716 0.09857
##                     nb_phylum   0.013039 0.11419  0.12
##  phylum_type        (Intercept) 0.234843 0.48461
##                     nb_phylum   0.030151 0.17364  0.55
##  empo_3             nb_phylum   0.014234 0.11931
## Number of obs: 33714, groups:
## empo_3:phylum_type, 911; empo_3:PI, 117; phylum_type, 84; empo_3, 17
##
## Fixed effects:
##             Estimate Std. Error z value Pr(>|z|)
## (Intercept)  0.30488    0.05807   5.250 1.52e-07 ***
## nb_phylum    0.27186    0.04288   6.341 2.29e-10 ***
## ---
## Signif. codes:  0 '***' 0.001 '**' 0.01 '*' 0.05 '.' 0.1 ' ' 1
```

nb__class=class:phylum, nb__phylum=number of non-focal phyla, phylum__type=Lineage, empo_3=Environment, PI=laboratory that submitted the data.

### 2. ASV based analysis

#### 2.1. ASV:Genus

```
## Generalized linear mixed model fit by maximum likelihood (Laplace
##   Approximation) [glmerMod]
##  Family: poisson  ( log )
## Formula:
## nb_ASV ~ shanon + (shanon | genus_type/empo_3) + (shanon | empo_3:PI) +
##     (shanon - 1 | sample)
##    Data: datsc1
## Control: glmerControl(optimizer = "bobyqa", optCtrl = list(maxfun = 1e+05))
##
##       AIC       BIC    logLik  deviance  df.resid
##  290622.3  290735.9 -145299.1  290598.3     95593
##
## Scaled residuals:
##     Min      1Q  Median      3Q     Max
## -3.6323 -0.3883 -0.1398  0.2125 12.2095
##
## Random effects:
##  Groups            Name        Variance Std.Dev. Corr
##  empo_3:genus_type (Intercept) 0.056752 0.23823
##                    shanon      0.022489 0.14996  0.16
##  sample            shanon      0.002968 0.05448
##  genus_type        (Intercept) 0.069456 0.26355
##                    shanon      0.006445 0.08028  0.70
##  empo_3:PI         (Intercept) 0.017813 0.13347
##                    shanon      0.007216 0.08495  0.20
## Number of obs: 95605, groups:
## empo_3:genus_type, 7857; sample, 1995; genus_type, 1128; empo_3:PI, 117
##
## Fixed effects:
##             Estimate Std. Error z value Pr(>|z|)
## (Intercept)  0.29051    0.01787   16.26  < 2e-16 ***
## shanon       0.05508    0.01272    4.33 1.49e-05 ***
## ---
## Signif. codes:  0 '***' 0.001 '**' 0.01 '*' 0.05 '.' 0.1 ' ' 1
```

#### 2.2. ASV:Family

```
## Generalized linear mixed model fit by maximum likelihood (Laplace
##   Approximation) [glmerMod]
##  Family: poisson  ( log )
## Formula:
## nb_ASV ~ shanon + (shanon | famil_type/empo_3) + (shanon | empo_3:PI) +
##     (shanon - 1 | sample) + (1 | obs)
##    Data: datsc2
```

```
## Control: glmerControl(optimizer = "bobyqa", optCtrl = list(maxfun = 1e+05))
##
##      AIC      BIC   logLik deviance df.resid
##  442753.1  442878.2 -221363.5  442727.1   112008
##
## Scaled residuals:
##     Min      1Q  Median      3Q     Max
## -2.6327 -0.4682 -0.1423  0.3132  3.6994
##
## Random effects:
##  Groups            Name        Variance Std.Dev. Corr
##  obs               (Intercept) 0.12959  0.3600
##  empo_3:famil_type (Intercept) 0.13062  0.3614
##                    shanon      0.07207  0.2685   0.01
##  sample            shanon      0.01786  0.1336
##  famil_type        (Intercept) 0.14374  0.3791
##                    shanon      0.03390  0.1841   0.59
##  empo_3:PI         (Intercept) 0.04177  0.2044
##                    shanon      0.02556  0.1599   -0.12
## Number of obs: 112021, groups:
## obs, 112021; empo_3:famil_type, 4833; sample, 1999; famil_type, 458; empo_3:PI, 117
##
## Fixed effects:
##             Estimate Std. Error z value Pr(>|z|)
## (Intercept)  0.50753    0.02995  16.949  < 2e-16 ***
## shanon       0.14766    0.02275   6.491 8.51e-11 ***
## ---
## Signif. codes:  0 '***' 0.001 '**' 0.01 '*' 0.05 '.' 0.1 ' ' 1
```

##### 2.3. ASV:Order

```
## Generalized linear mixed model fit by maximum likelihood (Laplace
##   Approximation) [glmerMod]
##  Family: poisson  ( log )
## Formula:
## nb_ASV ~ shanon + (shanon | order_type/empo_3) + (shanon | empo_3:PI) +
##     (shanon - 1 | sample) + (1 | obs)
##    Data: datsc2
## Control: glmerControl(optimizer = "bobyqa")
##
##      AIC      BIC   logLik deviance df.resid
##  486216.8  486341.1 -243095.4  486190.8   105181
##
## Scaled residuals:
##     Min      1Q  Median      3Q     Max
## -2.7884 -0.4696 -0.1030  0.2985  6.2226
##
## Random effects:
##  Groups            Name        Variance Std.Dev. Corr
##  obs               (Intercept) 0.19691  0.4437
##  empo_3:order_type (Intercept) 0.20007  0.4473
##                    shanon      0.17358  0.4166   -0.23
##  sample            shanon      0.04096  0.2024
```

```
##  order_type        (Intercept) 0.29459  0.5428
##                    shanon      0.11600  0.3406   0.33
##  empo_3:PI         (Intercept) 0.11781  0.3432
##                    shanon      0.06648  0.2578   0.09
## Number of obs: 105194, groups:
## obs, 105194; empo_3:order_type, 3781; sample, 2000; order_type, 363; empo_3:PI, 117
##
## Fixed effects:
##             Estimate Std. Error z value Pr(>|z|)
## (Intercept)  0.60375    0.04820  12.527   <2e-16 ***
## shanon       0.37796    0.03832   9.864   <2e-16 ***
## ---
## Signif. codes:  0 '***' 0.001 '**' 0.01 '*' 0.05 '.' 0.1 ' ' 1
```

##### 2.4. ASV:Class

```
## Generalized linear mixed model fit by maximum likelihood (Laplace
##   Approximation) [glmerMod]
##  Family: poisson  ( log )
## Formula:
## nb_ASV ~ shanon + (shanon | clas_type/empo_3) + (shanon | empo_3:PI) +
##     (shanon - 1 | sample) + (1 | obs)
##    Data: datsc2
## Control: glmerControl(optimizer = "bobyqa", optCtrl = list(maxfun = 1e+05))
##
##       AIC       BIC    logLik  deviance  df.resid
##  396695.5  396815.5 -198334.7  396669.5     75368
##
## Scaled residuals:
##     Min      1Q  Median      3Q     Max
## -2.5231 -0.4443 -0.0711  0.2558  8.3789
##
## Random effects:
##  Groups           Name        Variance Std.Dev. Corr
##  obs              (Intercept) 0.25714  0.5071
##  empo_3:clas_type (Intercept) 0.22980  0.4794
##                   shanon      0.21214  0.4606   -0.19
##  sample           shanon      0.06879  0.2623
##  clas_type        (Intercept) 0.50950  0.7138
##                   shanon      0.13654  0.3695   0.32
##  empo_3:PI        (Intercept) 0.19611  0.4428
##                   shanon      0.10615  0.3258   0.05
## Number of obs: 75381, groups:
## obs, 75381; empo_3:clas_type, 2383; sample, 2000; clas_type, 230; empo_3:PI, 117
##
## Fixed effects:
##             Estimate Std. Error z value Pr(>|z|)
## (Intercept)  0.63017    0.06863   9.182  < 2e-16 ***
## shanon       0.39836    0.04996   7.973 1.54e-15 ***
## ---
## Signif. codes:  0 '***' 0.001 '**' 0.01 '*' 0.05 '.' 0.1 ' ' 1
```

##### 2.5. ASV:Phylum

```
## Generalized linear mixed model fit by maximum likelihood (Laplace
##   Approximation) [glmerMod]
##  Family: poisson  ( log )
## Formula:
## nb_ASV ~ shanon + (shanon | phylum_type/empo_3) + (shanon | empo_3:PI) +
##     (shanon - 1 | empo_3) + (shanon - 1 | sample) + (1 | obs)
##    Data: datsc2
## Control: glmerControl(optimizer = "bobyqa", optCtrl = list(maxfun = 1e+05))
##
##       AIC       BIC    logLik  deviance  df.resid
##  211526.8  211644.8 -105749.4  211498.8     33700
##
## Scaled residuals:
##     Min      1Q  Median      3Q     Max
## -2.3449 -0.3817 -0.0319  0.1921  3.1857
##
## Random effects:
##  Groups             Name        Variance Std.Dev. Corr
##  obs                (Intercept) 0.31503  0.5613
##  sample             shanon      0.14296  0.3781
##  empo_3:phylum_type (Intercept) 0.28560  0.5344
##                     shanon      0.25029  0.5003   -0.27
##  empo_3:PI          (Intercept) 0.34088  0.5838
##                     shanon      0.24545  0.4954   -0.11
##  phylum_type        (Intercept) 1.13499  1.0654
##                     shanon      0.10031  0.3167   0.32
##  empo_3             shanon      0.02871  0.1694
## Number of obs: 33714, groups:
## obs, 33714; sample, 2000; empo_3:phylum_type, 911; empo_3:PI, 117; phylum_type, 84; empo_3, 17
##
## Fixed effects:
##             Estimate Std. Error z value Pr(>|z|)
## (Intercept)   0.5874     0.1360   4.320 1.56e-05 ***
## shanon        0.3187     0.0882   3.614 0.000302 ***
## ---
## Signif. codes:  0 '***' 0.001 '**' 0.01 '*' 0.05 '.' 0.1 ' ' 1
```

shanon=Shannon diversity of non focal lineages

#### 3. DBD is strongest in low-diversity environments

##### 3.1. Family:Order

```
## Generalized linear mixed model fit by maximum likelihood (Laplace
##   Approximation) [glmerMod]
##  Family: poisson  ( log )
## Formula: nb_family ~ nb_order * empo_3 + (nb_order | order_type) + (1 |
##     PI)
##    Data: datsc
## Control: glmerControl(optimizer = "bobyqa", optCtrl = list(maxfun = 1e+05))
##
```

```
##       AIC       BIC    logLik  deviance  df.resid
##  266705.4  267068.8 -133314.7  266629.4    105156
##
## Scaled residuals:
##     Min      1Q  Median      3Q     Max
## -2.7214 -0.1772 -0.0173  0.0831  6.7579
##
## Random effects:
##  Groups     Name        Variance Std.Dev. Corr
##  order_type (Intercept) 0.08678  0.2946
##             nb_order    0.01197  0.1094   0.45
##  PI         (Intercept) 0.00624  0.0790
## Number of obs: 105194, groups:  order_type, 363; PI, 61
##
## Fixed effects:
##                                    Estimate Std. Error z value Pr(>|z|)
## (Intercept)                         0.28087    0.02779  10.105  < 2e-16 ***
## nb_order                            0.13404    0.02016   6.648 2.97e-11 ***
## empo_3Animal corpus                 0.46962    0.10479   4.481 7.42e-06 ***
## empo_3Animal distal gut             0.30468    0.10115   3.012 0.002594 **
## empo_3Animal proximal gut           0.37860    0.09115   4.154 3.27e-05 ***
## empo_3Animal secretion              0.09225    0.03392   2.720 0.006536 **
## empo_3Animal surface                0.12120    0.02693   4.500 6.79e-06 ***
## empo_3Hypersaline (saline)         -0.10911    0.09228  -1.182 0.237064
## empo_3Plant corpus                  0.19482    0.03569   5.459 4.79e-08 ***
## empo_3Plant rhizosphere            -0.08185    0.02477  -3.304 0.000954 ***
## empo_3Plant surface                -0.01899    0.05932  -0.320 0.748923
## empo_3Sediment (non-saline)        -0.20406    0.02320  -8.797  < 2e-16 ***
## empo_3Sediment (saline)            -0.20619    0.02398  -8.597  < 2e-16 ***
## empo_3Soil (non-saline)            -0.09221    0.02214  -4.164 3.12e-05 ***
## empo_3Surface (non-saline)         -0.05198    0.02204  -2.358 0.018377 *
## empo_3Surface (saline)             -0.17916    0.05188  -3.453 0.000553 ***
## empo_3Water (non-saline)           -0.15775    0.02111  -7.474 7.79e-14 ***
## empo_3Water (saline)               -0.21450    0.02908  -7.377 1.62e-13 ***
## nb_order:empo_3Animal corpus        0.50958    0.07198   7.080 1.45e-12 ***
## nb_order:empo_3Animal distal gut    0.23727    0.07222   3.285 0.001019 **
## nb_order:empo_3Animal proximal gut  0.26375    0.06295   4.190 2.80e-05 ***
## nb_order:empo_3Animal secretion     0.15239    0.03171   4.806 1.54e-06 ***
## nb_order:empo_3Animal surface       0.12136    0.02909   4.172 3.02e-05 ***
## nb_order:empo_3Hypersaline (saline)  0.10875    0.10081   1.079 0.280678
## nb_order:empo_3Plant corpus         0.35619    0.03506  10.160  < 2e-16 ***
## nb_order:empo_3Plant rhizosphere   -0.15332    0.02421  -6.334 2.39e-10 ***
## nb_order:empo_3Plant surface        0.05029    0.03689   1.363 0.172749
## nb_order:empo_3Sediment (non-saline) -0.11854    0.02391  -4.958 7.14e-07 ***
## nb_order:empo_3Sediment (saline)   -0.03690    0.02397  -1.540 0.123657
## nb_order:empo_3Soil (non-saline)   -0.12247    0.02296  -5.335 9.56e-08 ***
## nb_order:empo_3Surface (non-saline)  -0.03367    0.02665  -1.263 0.206472
## nb_order:empo_3Surface (saline)    -0.03563    0.02335  -1.526 0.127049
## nb_order:empo_3Water (non-saline)  -0.09724    0.02315  -4.200 2.66e-05 ***
## nb_order:empo_3Water (saline)      -0.06915    0.02919  -2.369 0.017848 *
## ---
## Signif. codes:  0 '***' 0.001 '**' 0.01 '*' 0.05 '.' 0.1 ' ' 1

##
```

```
## Correlation matrix not shown by default, as p = 34 > 12.
## Use print(x, correlation=TRUE)  or
##     vcov(x)        if you need it
```

```
## convergence code: 0
## Model failed to converge with max|grad| = 0.00393273 (tol = 0.001, component 1)
```

nb_family=family:order, nb_order=number of non-focal orders, order_type=Lineage, empo_3=Environment, PI=laboratory that submitted the data.

##### 3.2. Order:Class

```
## Generalized linear mixed model fit by maximum likelihood (Laplace
##   Approximation) [glmerMod]
##  Family: poisson  ( log )
## Formula: nb_order ~ nb_classe * empo_3 + (nb_classe | classe_type) + (1 |
##     PI)
##    Data: datsc
## Control: glmerControl(optimizer = "bobyqa", optCtrl = list(maxfun = 1e+05))
##
##      AIC      BIC   logLik deviance df.resid
## 190975.6 191326.4 -95449.8 190899.6    75343
##
## Scaled residuals:
##     Min      1Q  Median      3Q     Max
## -2.4132 -0.1695 -0.0178  0.0773  3.7208
##
## Random effects:
##  Groups      Name        Variance Std.Dev. Corr
##  classe_type (Intercept) 0.115704 0.34015
##              nb_classe   0.024655 0.15702  0.59
##  PI          (Intercept) 0.001917 0.04378
## Number of obs: 75381, groups:  classe_type, 230; PI, 61
##
## Fixed effects:
##                                        Estimate Std. Error z value Pr(>|z|)
## (Intercept)                           0.1788354  0.0355370   5.032 4.84e-07
## nb_classe                             0.1202961  0.0271894   4.424 9.67e-06
## empo_3Animal corpus                   0.3129449  0.1011805   3.093  0.00198
## empo_3Animal distal gut               0.5368501  0.1310345   4.097 4.19e-05
## empo_3Animal proximal gut             0.1279622  0.1130821   1.132  0.25781
## empo_3Animal secretion                0.0687260  0.0435475   1.578  0.11452
## empo_3Animal surface                  0.0593722  0.0360301   1.648  0.09938
## empo_3Hypersaline (saline)            0.0005382  0.0728933   0.007  0.99411
## empo_3Plant corpus                    0.0833159  0.0434269   1.919  0.05504
## empo_3Plant rhizosphere               0.0515817  0.0301697   1.710  0.08732
## empo_3Plant surface                   0.0891866  0.0593787   1.502  0.13310
## empo_3Sediment (non-saline)           0.0326726  0.0290694   1.124  0.26103
## empo_3Sediment (saline)              -0.0258649  0.0288895  -0.895  0.37062
## empo_3Soil (non-saline)              -0.0103278  0.0275593  -0.375  0.70785
## empo_3Surface (non-saline)            0.0546641  0.0282431   1.935  0.05293
## empo_3Surface (saline)                0.0178921  0.0428232   0.418  0.67608
## empo_3Water (non-saline)             -0.0136723  0.0266687  -0.513  0.60818
## empo_3Water (saline)                  0.0320635  0.0328420   0.976  0.32892
```

```
## nb_classe:empo_3Animal corpus          0.3679761  0.0689111   5.340 9.30e-08
## nb_classe:empo_3Animal distal gut      0.5279010  0.0948001   5.569 2.57e-08
## nb_classe:empo_3Animal proximal gut    0.2401076  0.0805145   2.982  0.00286
## nb_classe:empo_3Animal secretion       0.1582968  0.0394070   4.017 5.90e-05
## nb_classe:empo_3Animal surface         0.0860404  0.0363760   2.365  0.01801
## nb_classe:empo_3Hypersaline (saline)   0.0578305  0.0884959   0.653  0.51345
## nb_classe:empo_3Plant corpus           0.3087379  0.0391894   7.878 3.32e-15
## nb_classe:empo_3Plant rhizosphere     -0.1322135  0.0289678  -4.564 5.02e-06
## nb_classe:empo_3Plant surface         -0.0327739  0.0407867  -0.804  0.42166
## nb_classe:empo_3Sediment (non-saline) -0.1197560  0.0289740  -4.133 3.58e-05
## nb_classe:empo_3Sediment (saline)     -0.0562754  0.0283242  -1.987  0.04694
## nb_classe:empo_3Soil (non-saline)     -0.0634347  0.0282739  -2.244  0.02486
## nb_classe:empo_3Surface (non-saline)   0.0329599  0.0323289   1.020  0.30796
## nb_classe:empo_3Surface (saline)      -0.0448347  0.0277785  -1.614  0.10653
## nb_classe:empo_3Water (non-saline)    -0.0844399  0.0277276  -3.045  0.00232
## nb_classe:empo_3Water (saline)        -0.0921891  0.0313269  -2.943  0.00325
##
## (Intercept)                           ***
## nb_classe                             ***
## empo_3Animal corpus                   **
## empo_3Animal distal gut               ***
## empo_3Animal proximal gut
## empo_3Animal secretion
## empo_3Animal surface                  .
## empo_3Hypersaline (saline)
## empo_3Plant corpus                    .
## empo_3Plant rhizosphere               .
## empo_3Plant surface
## empo_3Sediment (non-saline)
## empo_3Sediment (saline)
## empo_3Soil (non-saline)
## empo_3Surface (non-saline)            .
## empo_3Surface (saline)
## empo_3Water (non-saline)
## empo_3Water (saline)
## nb_classe:empo_3Animal corpus         ***
## nb_classe:empo_3Animal distal gut     ***
## nb_classe:empo_3Animal proximal gut   **
## nb_classe:empo_3Animal secretion      ***
## nb_classe:empo_3Animal surface        *
## nb_classe:empo_3Hypersaline (saline)
## nb_classe:empo_3Plant corpus          ***
## nb_classe:empo_3Plant rhizosphere     ***
## nb_classe:empo_3Plant surface
## nb_classe:empo_3Sediment (non-saline) ***
## nb_classe:empo_3Sediment (saline)     *
## nb_classe:empo_3Soil (non-saline)     *
## nb_classe:empo_3Surface (non-saline)
## nb_classe:empo_3Surface (saline)
## nb_classe:empo_3Water (non-saline)    **
## nb_classe:empo_3Water (saline)        **
## ---
## Signif. codes:  0 '***' 0.001 '**' 0.01 '*' 0.05 '.' 0.1 ' ' 1
```

```
##
## Correlation matrix not shown by default, as p = 34 > 12.
## Use print(x, correlation=TRUE)  or
##     vcov(x)        if you need it
```

```
## convergence code: 0
## Model failed to converge with max|grad| = 0.00163613 (tol = 0.001, component 1)
```

nb_order=order:class, nb_classe=number of non-focal classes, classe_type=Lineage, empo_3=Environment, PI=laboratory that submitted the data.

##### 3.3. Class:Phylum

```
## Generalized linear mixed model fit by maximum likelihood (Laplace
##   Approximation) [glmerMod]
##  Family: poisson  ( log )
## Formula: nb_class ~ nb_phylum * empo_3 + (nb_phylum | phylum_type) + (1 |
##     PI)
##    Data: datsc
## Control: glmerControl(optimizer = "bobyqa", optCtrl = list(maxfun = 1e+05))
##
##      AIC      BIC   logLik deviance df.resid
## 101518.6 101838.8 -50721.3 101442.6    33676
##
## Scaled residuals:
##     Min      1Q  Median      3Q     Max
## -2.6499 -0.3520 -0.0525  0.2605  4.3900
##
## Random effects:
##  Groups      Name        Variance Std.Dev. Corr
##  phylum_type (Intercept) 0.23697  0.48680
##              nb_phylum   0.03750  0.19364  0.42
##  PI          (Intercept) 0.00701  0.08373
## Number of obs: 33714, groups:  phylum_type, 84; PI, 61
##
## Fixed effects:
##                                     Estimate Std. Error z value Pr(>|z|)
## (Intercept)                          0.28661    0.06589   4.350 1.36e-05 ***
## nb_phylum                            0.33731    0.04234   7.968 1.62e-15 ***
## empo_3Animal corpus                  0.25901    0.10148   2.552 0.010698 *
## empo_3Animal distal gut              0.18777    0.11532   1.628 0.103483
## empo_3Animal proximal gut            0.16148    0.11773   1.372 0.170165
## empo_3Animal secretion               0.03595    0.05218   0.689 0.490780
## empo_3Animal surface                 0.10572    0.04419   2.392 0.016748 *
## empo_3Hypersaline (saline)          -0.12403    0.06233  -1.990 0.046581 *
## empo_3Plant corpus                   0.37243    0.05786   6.437 1.22e-10 ***
## empo_3Plant rhizosphere              0.10504    0.03769   2.787 0.005319 **
## empo_3Plant surface                  0.22745    0.08505   2.674 0.007487 **
## empo_3Sediment (non-saline)          0.13989    0.03634   3.849 0.000119 ***
## empo_3Sediment (saline)             -0.06715    0.03792  -1.771 0.076606 .
## empo_3Soil (non-saline)              0.09999    0.03523   2.838 0.004539 **
## empo_3Surface (non-saline)           0.11125    0.03711   2.998 0.002718 **
## empo_3Surface (saline)              -0.07392    0.06465  -1.144 0.252824
## empo_3Water (non-saline)            -0.04040    0.03464  -1.166 0.243522
```

```
## empo_3Water (saline)                  -0.14358    0.03964  -3.622 0.000292 ***
## nb_phylum:empo_3Animal corpus          0.36484    0.07850   4.647 3.36e-06 ***
## nb_phylum:empo_3Animal distal gut      0.18855    0.09250   2.038 0.041520 *
## nb_phylum:empo_3Animal proximal gut    0.18888    0.09837   1.920 0.054862 .
## nb_phylum:empo_3Animal secretion       0.06901    0.05120   1.348 0.177706
## nb_phylum:empo_3Animal surface         0.11961    0.05051   2.368 0.017892 *
## nb_phylum:empo_3Hypersaline (saline)  -0.14872    0.08425  -1.765 0.077524 .
## nb_phylum:empo_3Plant corpus           0.50461    0.05625   8.970  < 2e-16 ***
## nb_phylum:empo_3Plant rhizosphere     -0.25721    0.04138  -6.216 5.09e-10 ***
## nb_phylum:empo_3Plant surface          0.06277    0.05969   1.051 0.293046
## nb_phylum:empo_3Sediment (non-saline) -0.36355    0.03785  -9.605  < 2e-16 ***
## nb_phylum:empo_3Sediment (saline)     -0.27209    0.03771  -7.215 5.38e-13 ***
## nb_phylum:empo_3Soil (non-saline)     -0.25006    0.03936  -6.353 2.11e-10 ***
## nb_phylum:empo_3Surface (non-saline)  -0.04118    0.04476  -0.920 0.357563
## nb_phylum:empo_3Surface (saline)      -0.19153    0.03754  -5.102 3.35e-07 ***
## nb_phylum:empo_3Water (non-saline)    -0.20028    0.03837  -5.220 1.79e-07 ***
## nb_phylum:empo_3Water (saline)        -0.21296    0.04117  -5.173 2.30e-07 ***
## ---
## Signif. codes:  0 '***' 0.001 '**' 0.01 '*' 0.05 '.' 0.1 ' ' 1
##
## Correlation matrix not shown by default, as p = 34 > 12.
## Use print(x, correlation=TRUE)  or
##     vcov(x)        if you need it
```

nb_class=class:phylum, nb_phylum=number of non-focal phyla, phylum_type=Lineage, empo_3=Environment, PI=laboratory that submitted the data.

### 4. Abiotic factors analysis

#### 4.1. ASV:Genus

```
## Generalized linear mixed model fit by maximum likelihood (Laplace
##   Approximation) [glmerMod]
##  Family: poisson  ( log )
## Formula: nb_ASV ~ nb_genus + temperature_deg_c + nb_genus:temperature_deg_c +
##     nb_genus:latitude_deg + nb_genus:elevation_m + (1 | genus_type/empo_3) +
##     (1 | empo_3:PI) + (1 | sample)
##    Data: datsc
## Control: glmerControl(optimizer = "bobyqa")
##
##      AIC      BIC   logLik deviance df.resid
##  27838.5  27909.7 -13909.3  27818.5     9112
##
## Scaled residuals:
##     Min      1Q  Median      3Q     Max
## -3.3363 -0.3680 -0.1554  0.1999  8.9640
##
## Random effects:
##  Groups            Name        Variance Std.Dev.
##  empo_3:genus_type (Intercept) 0.087128 0.29517
##  genus_type        (Intercept) 0.091049 0.30174
##  sample            (Intercept) 0.004665 0.06830
```

```
##  empo_3:PI          (Intercept) 0.003740 0.06116
## Number of obs: 9122, groups:
## empo_3:genus_type, 1740; genus_type, 651; sample, 192; empo_3:PI, 16
##
## Fixed effects:
##                            Estimate Std. Error z value Pr(>|z|)
## (Intercept)                 0.32688    0.02773  11.788  < 2e-16 ***
## nb_genus                    0.12837    0.01330   9.652  < 2e-16 ***
## temperature_deg_c           0.03995    0.01416   2.821  0.00479 **
## nb_genus:temperature_deg_c  0.04340    0.01387   3.129  0.00175 **
## nb_genus:latitude_deg       0.03112    0.01350   2.305  0.02119 *
## nb_genus:elevation_m       -0.03137    0.01430  -2.193  0.02829 *
## ---
## Signif. codes:  0 '***' 0.001 '**' 0.01 '*' 0.05 '.' 0.1 ' ' 1
```

nb_ASV=ASV:family, nb_genus=number of non-focal genera, genus_type=Lineage, empo_3=Environment, PI=laboratory that submitted the data, sample=EMP sample ID, temperature_deg_c=temperature, latitude_deg=latitude, elevation_m=elevation.

#### 4.2. Genus:Family

```
## Generalized linear mixed model fit by maximum likelihood (Laplace
##   Approximation) [glmerMod]
##  Family: poisson  ( log )
## Formula:
## nb_genus ~ nb_family + temperature_deg_c + ph + (1 | family_type/empo_3)
##    Data: datsc
## Control: glmerControl(optimizer = "bobyqa")
##
##      AIC      BIC   logLik deviance df.resid
##  31081.0  31125.7 -15534.5  31069.0    12624
##
## Scaled residuals:
##     Min      1Q  Median      3Q     Max
## -1.6402 -0.2080 -0.0755  0.0507  5.2865
##
## Random effects:
##  Groups             Name        Variance Std.Dev.
##  empo_3:family_type (Intercept) 0.02926  0.1711
##  family_type        (Intercept) 0.05404  0.2325
## Number of obs: 12630, groups:  empo_3:family_type, 1582; family_type, 389
##
## Fixed effects:
##                    Estimate Std. Error z value Pr(>|z|)
## (Intercept)        0.175100   0.017917   9.773  < 2e-16 ***
## nb_family          0.094498   0.008669  10.901  < 2e-16 ***
## temperature_deg_c  0.026323   0.008768   3.002  0.00268 **
## ph                -0.041609   0.009184  -4.531 5.88e-06 ***
## ---
## Signif. codes:  0 '***' 0.001 '**' 0.01 '*' 0.05 '.' 0.1 ' ' 1
```

nb_genus=genus:family, nb_family=number of non-focal families, family_type=Lineage, empo_3=Environment, temperature_deg_c=temperature, latitude_deg=latitude, elevation_m=elevation.

##### 4.3. Family:Order

```
## Generalized linear mixed model fit by maximum likelihood (Laplace
##   Approximation) [glmerMod]
##  Family: poisson  ( log )
## Formula: nb_family ~ nb_order + (1 | order_type/empo_3) + (1 | empo_3:PI)
##    Data: datsc
## Control: glmerControl(optimizer = "bobyqa")
##
##      AIC      BIC   logLik deviance df.resid
##  32598.6  32636.0 -16294.3  32588.6    12985
##
## Scaled residuals:
##     Min      1Q  Median      3Q     Max
## -1.6667 -0.1693 -0.0398  0.0872  3.9125
##
## Random effects:
##  Groups            Name        Variance Std.Dev.
##  empo_3:order_type (Intercept) 0.02992  0.1730
##  order_type        (Intercept) 0.09716  0.3117
##  empo_3:PI         (Intercept) 0.01017  0.1008
## Number of obs: 12990, groups:
## empo_3:order_type, 1360; order_type, 323; empo_3:PI, 16
##
## Fixed effects:
##             Estimate Std. Error z value Pr(>|z|)
## (Intercept)  0.17547    0.03567   4.919 8.69e-07 ***
## nb_order     0.13134    0.01092  12.028  < 2e-16 ***
## ---
## Signif. codes:  0 '***' 0.001 '**' 0.01 '*' 0.05 '.' 0.1 ' ' 1
```

nb_family=family:order, nb_order=number of non-focal orders, order_type=Lineage, empo_3=Environment, PI=laboratory that submitted the data, temperature_deg_c=temperature, latitude_deg=latitude, elevation_m=elevation.

##### 4.4. Order:Class

```
## Generalized linear mixed model fit by maximum likelihood (Laplace
##   Approximation) [glmerMod]
##  Family: poisson  ( log )
## Formula:
## nb_order ~ nb_classe + nb_classe:temperature_deg_c + nb_classe:latitude_deg +
##     (1 | classe_type/empo_3)
##    Data: datsc
## Control: glmerControl(optimizer = "bobyqa")
##
##      AIC      BIC   logLik deviance df.resid
##  24298.7  24341.6 -12143.3  24286.7     9453
##
## Scaled residuals:
##     Min      1Q  Median      3Q     Max
## -1.9938 -0.1883 -0.0498  0.1533  3.3081
##
## Random effects:
```

```
##  Groups             Name        Variance Std.Dev.
##  empo_3:classe_type (Intercept) 0.03287  0.1813
##  classe_type        (Intercept) 0.15383  0.3922
## Number of obs: 9459, groups:  empo_3:classe_type, 941; classe_type, 206
##
## Fixed effects:
##                             Estimate Std. Error z value Pr(>|z|)
## (Intercept)                 0.151710   0.033274   4.559 5.13e-06 ***
## nb_classe                   0.184448   0.009933  18.570  < 2e-16 ***
## nb_classe:temperature_deg_c 0.031932   0.009550   3.344 0.000827 ***
## nb_classe:latitude_deg      0.023453   0.008431   2.782 0.005403 **
## ---
## Signif. codes:  0 '***' 0.001 '**' 0.01 '*' 0.05 '.' 0.1 ' ' 1
```

nb_order=order:class, nb_classe=number of non-focal classes, classe_type=Lineage, empo_3=Environment, temperature_deg_c=temperature, latitude_deg=latitude, elevation_m=elevation.

##### 4.5. Class:Phylum

```
## Generalized linear mixed model fit by maximum likelihood (Laplace
##   Approximation) [glmerMod]
##  Family: poisson  ( log )
## Formula:
## nb_class ~ nb_phylum + nb_phylum:temperature_deg_c + nb_phylum:latitude_deg +
##     nb_phylum:ph + (1 | phylum_type/empo_3)
##    Data: datsc
## Control: glmerControl(optimizer = "bobyqa")
##
##      AIC      BIC   logLik deviance df.resid
##  12482.4  12526.7  -6234.2  12468.4     4101
##
## Scaled residuals:
##     Min      1Q  Median      3Q     Max
## -1.5567 -0.3108 -0.0519  0.2560  3.7076
##
## Random effects:
##  Groups             Name        Variance Std.Dev.
##  empo_3:phylum_type (Intercept) 0.04314  0.2077
##  phylum_type        (Intercept) 0.31013  0.5569
## Number of obs: 4108, groups:  empo_3:phylum_type, 367; phylum_type, 78
##
## Fixed effects:
##                             Estimate Std. Error z value Pr(>|z|)
## (Intercept)                  0.25091    0.07113   3.527  0.00042 ***
## nb_phylum                    0.23662    0.01191  19.864  < 2e-16 ***
## nb_phylum:temperature_deg_c  0.05921    0.01394   4.249 2.15e-05 ***
## nb_phylum:latitude_deg       0.03013    0.01151   2.618  0.00884 **
## nb_phylum:ph                 0.01263    0.01090   1.159  0.24661
## ---
## Signif. codes:  0 '***' 0.001 '**' 0.01 '*' 0.05 '.' 0.1 ' ' 1
```

nb_class=class:phylum, nb_phylum=number of non-focal phyla, phylum_type=Lineage, empo_3=Environment, temperature_deg_c=temperature, latitude_deg=latitude, elevation_m=elevation.

### 5. DBD in resident genera versus non-resident genera

#### 5.1. Animal cluster

```
## Generalized linear mixed model fit by maximum likelihood (Laplace
##   Approximation) [glmerMod]
##  Family: poisson  ( log )
## Formula: nb_ASV ~ nb_genus * status + (nb_genus | genus_type/empo_3) +
##     (nb_genus | empo_3:PI) + (nb_genus - 1 | sample)
##    Data: datsc1
## Control: glmerControl(optimizer = "bobyqa")
##
##      AIC      BIC   logLik deviance df.resid
##  98718.5  98853.1 -49343.2  98686.5    33295
##
## Scaled residuals:
##     Min      1Q  Median      3Q     Max
## -3.0576 -0.3761 -0.1504  0.2038  8.2981
##
## Random effects:
##  Groups            Name        Variance Std.Dev. Corr
##  empo_3:genus_type (Intercept) 0.039726 0.19931
##                    nb_genus    0.008227 0.09070  0.17
##  genus_type        (Intercept) 0.063582 0.25216
##                    nb_genus    0.007117 0.08436  0.88
##  sample            nb_genus    0.004299 0.06557
##  empo_3:PI         (Intercept) 0.028807 0.16973
##                    nb_genus    0.007302 0.08545  0.11
## Number of obs: 33311, groups:
## empo_3:genus_type, 3196; genus_type, 850; sample, 795; empo_3:PI, 35
##
## Fixed effects:
##                         Estimate Std. Error z value Pr(>|z|)
## (Intercept)              0.25300    0.03799   6.659 2.75e-11 ***
## nb_genus                 0.01619    0.02505   0.646    0.518
## statusMigrant            0.04152    0.03492   1.189    0.234
## statusNative             0.15219    0.03129   4.863 1.15e-06 ***
## nb_genus:statusMigrant -0.01142    0.02018  -0.566    0.571
## nb_genus:statusNative   0.11791    0.01661   7.100 1.25e-12 ***
## ---
## Signif. codes:  0 '***' 0.001 '**' 0.01 '*' 0.05 '.' 0.1 ' ' 1
```

nb_ASV=ASV:genus, nb_genus=number of non-focal native genera, genus_type=Lineage, empo_3=Environment, PI=laboratory that submitted the data, sample=EMP sample ID, status=Migrant/Native (Resident)/Generalist.

#### 5.2. Saline cluster

```
## Generalized linear mixed model fit by maximum likelihood (Laplace
##   Approximation) [glmerMod]
##  Family: poisson  ( log )
## Formula: nb_ASV ~ nb_genus * status + (nb_genus | genus_type/empo_3) +
##     (nb_genus | empo_3:PI) + (nb_genus - 1 | sample)
```

```
##    Data: datsc1
## Control: glmerControl(optimizer = "bobyqa")
##
##      AIC      BIC   logLik deviance df.resid
##  57099.8  57225.9 -28533.9  57067.8    19486
##
## Scaled residuals:
##     Min      1Q  Median      3Q     Max
## -3.3816 -0.3677 -0.1566  0.1609  8.1450
##
## Random effects:
##  Groups            Name        Variance Std.Dev. Corr
##  empo_3:genus_type (Intercept) 0.037468 0.19357
##                    nb_genus    0.005197 0.07209  -0.15
##  genus_type        (Intercept) 0.080367 0.28349
##                    nb_genus    0.006384 0.07990  0.36
##  sample            nb_genus    0.004304 0.06561
##  empo_3:PI         (Intercept) 0.013523 0.11629
##                    nb_genus    0.004867 0.06976  -0.41
## Number of obs: 19502, groups:
## empo_3:genus_type, 1998; genus_type, 788; sample, 498; empo_3:PI, 26
##
## Fixed effects:
##                         Estimate Std. Error z value Pr(>|z|)
## (Intercept)             0.270074   0.033510   8.060 7.65e-16 ***
## nb_genus                0.006229   0.022320   0.279 0.780173
## statusMigrant          -0.006777   0.038090  -0.178 0.858789
## statusNative           -0.005741   0.043218  -0.133 0.894323
## nb_genus:statusMigrant -0.070906   0.022200  -3.194 0.001404 **
## nb_genus:statusNative   0.071725   0.021632   3.316 0.000914 ***
## ---
## Signif. codes:  0 '***' 0.001 '**' 0.01 '*' 0.05 '.' 0.1 ' ' 1
```

nb_ASV=ASV:genus, nb_genus=number of non-focal native genera, genus_type=Lineage, empo_3=Environment, PI=laboratory that submitted the data, sample=EMP sample ID, status=Migrant/Native (Resident)/Generalist.

#### 5.3. Non saline cluster

```
## Generalized linear mixed model fit by maximum likelihood (Laplace
##   Approximation) [glmerMod]
##  Family: poisson  ( log )
## Formula: nb_ASV ~ nb_genus * status + (nb_genus | genus_type/empo_3) +
##     (nb_genus | empo_3:PI) + (nb_genus - 1 | sample)
##    Data: datsc1
## Control: glmerControl(optimizer = "bobyqa")
##
##      AIC      BIC   logLik deviance df.resid
## 131340.7 131479.0 -65654.3 131308.7    42063
##
## Scaled residuals:
##     Min      1Q  Median      3Q     Max
## -3.4818 -0.3928 -0.1428  0.2505 10.2862
##
```

```
## Random effects:
##  Groups            Name        Variance Std.Dev. Corr
##  empo_3:genus_type (Intercept) 0.042733 0.20672
##                    nb_genus    0.029340 0.17129  0.04
##  genus_type        (Intercept) 0.097078 0.31157
##                    nb_genus    0.008959 0.09465  0.70
##  sample            nb_genus    0.001827 0.04274
##  empo_3:PI         (Intercept) 0.011668 0.10802
##                    nb_genus    0.013691 0.11701  0.04
## Number of obs: 42079, groups:
## empo_3:genus_type, 2611; genus_type, 870; sample, 637; empo_3:PI, 55
##
## Fixed effects:
##                         Estimate Std. Error z value Pr(>|z|)
## (Intercept)             0.251500   0.028175   8.926  < 2e-16 ***
## nb_genus                0.008664   0.025004   0.347  0.72896
## statusMigrant           0.020687   0.044937   0.460  0.64527
## statusNative            0.233262   0.035794   6.517 7.19e-11 ***
## nb_genus:statusMigrant -0.079900   0.028135  -2.840  0.00451 **
## nb_genus:statusNative   0.141488   0.019904   7.109 1.17e-12 ***
## ---
## Signif. codes:  0 '***' 0.001 '**' 0.01 '*' 0.05 '.' 0.1 ' ' 1
```

nb_ASV=ASV:genus, nb_genus=number of non-focal native genera, genus_type=Lineage, empo_3=Environment, PI=laboratory that submitted the data, sample=EMP sample ID, status=Migrant/Native (Resident)/Generalist.

### 6. Genome size analysis

```
## Generalized linear mixed model fit by maximum likelihood (Laplace
##   Approximation) [glmerMod]
##  Family: poisson  ( log )
## Formula:
## nb_ASV ~ nb_genus * size + (nb_genus | genus_type/empo_3) + (nb_genus |
##     empo_3:PI) + (nb_genus - 1 | sample)
##    Data: datsc1
## Control: glmerControl(optimizer = "bobyqa")
##
##       AIC       BIC    logLik  deviance  df.resid
##  240595.8  240725.6 -120283.9  240567.8     78199
##
## Scaled residuals:
##     Min      1Q  Median      3Q     Max
## -3.6998 -0.4020 -0.1466  0.2450 11.9407
##
## Random effects:
##  Groups            Name        Variance Std.Dev. Corr
##  empo_3:genus_type (Intercept) 0.059649 0.24423
##                    nb_genus    0.021223 0.14568  0.05
##  sample            nb_genus    0.002721 0.05216
##  genus_type        (Intercept) 0.074050 0.27212
##                    nb_genus    0.005595 0.07480  0.60
##  empo_3:PI         (Intercept) 0.017041 0.13054
```

```
##                    nb_genus    0.013100 0.11445  0.18
## Number of obs: 78213, groups:
## empo_3:genus_type, 5501; sample, 1993; genus_type, 576; empo_3:PI, 117
##
## Fixed effects:
##               Estimate Std. Error z value Pr(>|z|)
## (Intercept)   0.371754   0.020487  18.145  < 2e-16 ***
## nb_genus      0.113302   0.016313   6.945 3.77e-12 ***
## size          0.001268   0.014027   0.090   0.9280
## nb_genus:size 0.015908   0.006342   2.508   0.0121 *
## ---
## Signif. codes:  0 '***' 0.001 '**' 0.01 '*' 0.05 '.' 0.1 ' ' 1
```

nb_ASV=ASV:genus, nb_genus=number of non-focal genera, genus_type=Lineage, empo_3=Environment, PI=laboratory that submitted the data, sample=EMP sample ID, size=genome size.
