## Supplementary material for "Does diversity beget diversity in microbiomes?": File S2

**Table S1.** **Goodness of fit for the GLMMs**. We report the marginal (*R^2^*M) and conditional (*R^2^*C) coefficients of variation, which respectively include variance explained by fixed effects only, and by fixed and random effects. In the right-most two columns, the difference in AIC (∆AIC) and LRT *P-*values are reported comparing the full model to a model with no fixed effects other than the intercept. The top panel (1) shows GLMMs fitted on the EMP data, and the bottom (2) shows the soil data. Note that no abiotic or biotic factors were significant at the order:class level in soil.

1. EMP data

|  | ***R^2^* M** | | ***R^2^*C** | | **Comparison to null model** | |
| --- | --- | --- | --- | --- | --- | --- |
|  | **Lognormal** | **Trigamma** | **Lognormal** | **Trigamma** | **∆AIC** | ***P*-value** |
| **Taxonomy-based GLMMs (Table 1)** | | | | |  |  |
| **ASV :Genus** | 0.012 | 0.007 | 0.273 | 0.17 | 29 | 3.48e-08 |
| **Genus :Family** | 0.003 | 0.002 | 0.127 | 0.065 | 29 | 2.358e-08 |
| **Family :Order** | 0.019 | 0.011 | 0.212 | 0.118 | 23 | 4.229e-07 |
| **Order :Class** | 0.016 | 0.009 | 0.226 | 0.127 | 20 | 2.473e-06 |
| **Class :Phylum** | 0.082 | 0.058 | 0.464 | 0.33 | 23 | 5.618e-07 |
| **DBD variation across environments (Figure 3)** | | | | |  |  |
| **Family:order** | 0.015 | 0.008 | 0.158 | 0.081 | 948.53 | < 2.2e-16 |
| **Order:class** | 0.026 | 0.013 | 0.208 | 0.11 | 691.74 | < 2.2e-16 |
| **Class:phylum** | 0.074 | 0.046 | 0.383 | 0.24 | 1059.45 | < 2.2e-16 |
| **Abiotic factors (Table 4)** | | | | |  |  |
| **ASV :Genus** | 0.027 | 0.017 | 0.283 | 0.176 | 93 | < 2.2e-16 |
| **Genus :Family** | 0.014 | 0.007 | 0.137 | 0.072 | 133 | < 2.2e-16 |
| **Family :Order** | 0.023 | 0.012 | 0.2 | 0.11 | 140 | < 2.2e-16 |
| **Order :Class** | 0.046 | 0.027 | 0.281 | 0.165 | 374 | < 2.2e-16 |
| **Class :Phylum** | 0.068 | 0.048 | 0.46 | 0.326 | 411 | < 2.2e-16 |
| **DBD in resident versus non-resident genera (Figure 4)** | | | | |  |  |
| **animal** | 0.019 | 0.011 | 0.246 | 0.148 | 57 | 5.69e-13 |
| **saline** | 0.004 | 0.002 | 0.225 | 0.134 | 18 | 4.051e-05 |
| **Non saline** | 0.036 | 0.023 | 0.316 | 0.203 | 88 | < 2.2e-16 |
| **Genome size analysis (Figure 5)** | | | | |  |  |
| **ASV:Genus** | 0.018 | 0.012 | 0.293 | 0.187 | 42 | 2.009e-10 |

2. Soil data (Table 5)

|  | ***R^2^* M** | | ***R^2^* C** | | **Comparison to null model** | |
| --- | --- | --- | --- | --- | --- | --- |
|  | **Lognormal** | **Trigamma** | **Lognormal** | **Trigamma** | **∆AIC** | ***P*-value** |
| **1. GLMMs with abiotic variables** | | | | | | |
| **ASV :Genus** | 0.016 | 0.013 | 0.519 | 0.405 | 783.062 | <2.2e-16 |
| **Genus :Family** | 0.003 | 0.002 | 0.253 | 0.149 | 43 | 4.846e-11 |
| **Family :Order** | 0.000 | 0.000 | 0.26 | 0.153 | 4 | 0.01073 |
| **Class :Phylum** | 0.008 | 0.007 | 0.542 | 0.431 | 71 | < 2.2e-16 |
| **2.GLMMs with PCs** | | | | | | |
| **ASV :Genus** | 0.014 | 0.01 | 0.526 | 0.408 | 103 | < 2.2e-16 |
| **Genus :Family** | 0.003 | 0.002 | 0.252 | 0.149 | 23 | 2.067e-06 |
| **Family :Order** | 0.006 | 0.004 | 0.266 | 0.157 | 103 | < 2.2e-16 |
| **Class :Phylum** | 0.006 | 0.005 | 0.541 | 0.429 | 51 | 3.141e-12 |
