## Supplementary material for "Does diversity beget diversity in microbiomes?": File S3

### Full GLMMs output for neutral model synthetic data

Significant models (Likelihood-ratio test, p<0.05)

Naïma Madi

### 1. ZSM distribution data (model 1 and model 2)

#### 1.1. Basic GLMMs

Models with focal lineage diversity as a function of community diversity in genus:ASV level, with genus name (genus_type) and EMP sample id (sample) as random effects in model 1 (data generated without environment information, one distribution for all) :
model 1 = glmer(nb__ASV~nb__genus+(nb__genus|genus__type)+(1|sample))
Environment type (empo__3) was added as a random effect in model 2 (data generated with a different distribution for each environment):
model 2 = glmer(nb__ASV~nb__genus+(nb__genus|genus__type/empo__3)+(nb__genus|empo__3)+(1|sample))

##### 1.1.1. Model 1

Significative negative effect of community diversity on focal lineage diversity

```
## Generalized linear mixed model fit by maximum likelihood (Laplace
##   Approximation) [glmerMod]
##  Family: poisson  ( log )
## Formula: nb_ASV ~ nb_genus + (1 | genus_type)
##    Data: datsc1
## Control: glmerControl(optimizer = "bobyqa", optCtrl = list(maxfun = 1e+05))
##
##      AIC      BIC   logLik deviance df.resid
##  3220595  3220630 -1610294  3220589  1014686
##
## Scaled residuals:
##     Min      1Q  Median      3Q     Max
## -3.1977 -0.4064 -0.1064  0.3593  5.5833
##
## Random effects:
##  Groups     Name        Variance Std.Dev.
##  genus_type (Intercept) 0.4077   0.6385
## Number of obs: 1014689, groups:  genus_type, 1128
##
## Fixed effects:
##               Estimate Std. Error z value Pr(>|z|)
## (Intercept)  0.4433438  0.0189682  23.373   <2e-16 ***
## nb_genus    -0.0051824  0.0005284  -9.807   <2e-16 ***
## ---
## Signif. codes:  0 '***' 0.001 '**' 0.01 '*' 0.05 '.' 0.1 ' ' 1
```

```
## convergence code: 0
## Model is nearly unidentifiable: very large eigenvalue
##  - Rescale variables?
```

##### 1.1.2. Model 2

Null model (intercept only) is significant, meaning no significant effect of community diversity on the focal lineage diversity.

```
## Generalized linear mixed model fit by maximum likelihood (Laplace
##   Approximation) [glmerMod]
##  Family: poisson  ( log )
## Formula: nb_ASV ~ 1 + (1 | genus_type/empo_3) + (nb_genus | empo_3)
##    Data: datsc1
## Control: glmerControl(optimizer = "bobyqa", optCtrl = list(maxfun = 1e+05))
##
##      AIC      BIC   logLik deviance df.resid
##  3171080  3171151 -1585534  3171068  1001228
##
## Scaled residuals:
##     Min      1Q  Median      3Q     Max
## -3.4246 -0.4097 -0.1145  0.3470  5.2430
##
## Random effects:
##  Groups            Name        Variance  Std.Dev. Corr
##  empo_3:genus_type (Intercept) 2.682e-04 0.016377
##  genus_type        (Intercept) 4.012e-01 0.633416
##  empo_3            (Intercept) 7.434e-03 0.086223
##                    nb_genus    2.865e-06 0.001693 0.21
## Number of obs: 1001234, groups:
## empo_3:genus_type, 19085; genus_type, 1128; empo_3, 17
##
## Fixed effects:
##             Estimate Std. Error z value Pr(>|z|)
## (Intercept)  0.43044    0.02802   15.36   <2e-16 ***
## ---
## Signif. codes:  0 '***' 0.001 '**' 0.01 '*' 0.05 '.' 0.1 ' ' 1
## convergence code: 0
## Model failed to converge with max|grad| = 0.00397647 (tol = 0.001, component 1)
## Model is nearly unidentifiable: very large eigenvalue
##  - Rescale variables?
```

#### 1.2. Biome effect on DBD (model 2)

Model with focal lineage diversity as a function of the interaction between community diversity and environment type for model 2 (data generated with a different distribution for each biome). This model was ran for taxonomic ratios with significant DBD slope variation by environment in the EMP data
model = glmer(nb_class~nb_phylum*empo_3+(nb_phylum|phylum_type)+(1|sample))
model = glmer(nb_order~nb_class*empo_3+(nb_class|class_type)+(1|sample))
model = glmer(nb_family~nb_order*empo_3+(nb_order|order_type)+(1|sample))

##### 1.2.1. Class:Phylum

Null model (intercept only) is significant.

```
## Generalized linear mixed model fit by maximum likelihood (Laplace
##   Approximation) [glmerMod]
##  Family: poisson  ( log )
## Formula: nb_class ~ 1 + (1 | phylum_type)
##    Data: datsc1
## Control: glmerControl(optimizer = "bobyqa")
##
##      AIC      BIC   logLik deviance df.resid
## 134155.7 134173.4 -67075.8 134151.7    51981
##
## Scaled residuals:
##      Min       1Q   Median       3Q      Max
## -1.40666 -0.00089 -0.00057  0.00059  1.25225
##
## Random effects:
##  Groups      Name        Variance Std.Dev.
##  phylum_type (Intercept) 0.4431   0.6657
## Number of obs: 51983, groups:  phylum_type, 31
##
## Fixed effects:
##             Estimate Std. Error z value Pr(>|z|)
## (Intercept)   0.5018     0.1170   4.289 1.79e-05 ***
## ---
## Signif. codes:  0 '***' 0.001 '**' 0.01 '*' 0.05 '.' 0.1 ' ' 1
```

##### 1.2.2. Order:Class

Null model (intercept only) is significant.

```
## Generalized linear mixed model fit by maximum likelihood (Laplace
##   Approximation) [glmerMod]
##  Family: poisson  ( log )
## Formula: nb_order ~ 1 + (1 | class_type)
##    Data: datsc1
## Control: glmerControl(optimizer = "bobyqa", optCtrl = list(maxfun = 2e+05))
##
##       AIC       BIC    logLik  deviance  df.resid
##  284436.2  284455.6 -142216.1  284432.2    121228
##
## Scaled residuals:
##      Min       1Q   Median       3Q      Max
## -1.14082 -0.00057 -0.00049 -0.00049  2.22002
##
## Random effects:
##  Groups     Name        Variance Std.Dev.
##  class_type (Intercept) 0.2851   0.534
## Number of obs: 121230, groups:  class_type, 75
##
## Fixed effects:
##             Estimate Std. Error z value Pr(>|z|)
```

```
## (Intercept)  0.28183    0.06039   4.667 3.06e-06 ***
## ---
## Signif. codes:  0 '***' 0.001 '**' 0.01 '*' 0.05 '.' 0.1 ' ' 1
```

###### 1.2.3. Family:Order

Model with intercept varying by biome is significant.

```
## Generalized linear mixed model fit by maximum likelihood (Laplace
##   Approximation) [glmerMod]
##  Family: poisson  ( log )
## Formula: nb_family ~ empo_3 + (1 | order_type)
##    Data: datsc1
## Control: glmerControl(optimizer = "bobyqa", optCtrl = list(maxfun = 2e+05))
##
##       AIC       BIC    logLik  deviance  df.resid
##  531270.0  531455.3 -265617.0  531234.0    218252
##
## Scaled residuals:
##      Min       1Q   Median       3Q      Max
## -1.70252 -0.02264  0.00016  0.02217  1.77064
##
## Random effects:
##  Groups     Name        Variance Std.Dev.
##  order_type (Intercept) 0.3056   0.5528
## Number of obs: 218270, groups:  order_type, 141
##
## Fixed effects:
##                             Estimate Std. Error z value Pr(>|z|)
## (Intercept)                 0.320427   0.046181   6.939 3.96e-12 ***
## empo_3Animal corpus         0.029108   0.009535   3.053 0.002267 **
## empo_3Animal distal gut     0.007655   0.009601   0.797 0.425229
## empo_3Animal proximal gut   0.012015   0.009587   1.253 0.210130
## empo_3Animal secretion     -0.005246   0.009882  -0.531 0.595548
## empo_3Animal surface        0.035064   0.009337   3.755 0.000173 ***
## empo_3Hypersaline (saline)  0.028361   0.019865   1.428 0.153381
## empo_3Plant corpus          0.028393   0.009611   2.954 0.003133 **
## empo_3Plant rhizosphere     0.011224   0.009588   1.171 0.241787
## empo_3Plant surface        -0.010235   0.009655  -1.060 0.289144
## empo_3Sediment (non-saline)  0.019899   0.009566   2.080 0.037508 *
## empo_3Sediment (saline)     0.017615   0.009571   1.841 0.065683 .
## empo_3Soil (non-saline)     0.034402   0.009503   3.620 0.000295 ***
## empo_3Surface (non-saline)  0.019388   0.009564   2.027 0.042638 *
## empo_3Surface (saline)      0.005603   0.009783   0.573 0.566864
## empo_3Water (non-saline)    0.005154   0.009593   0.537 0.591066
## empo_3Water (saline)       -0.032116   0.009718  -3.305 0.000951 ***
## ---
## Signif. codes:  0 '***' 0.001 '**' 0.01 '*' 0.05 '.' 0.1 ' ' 1

##
## Correlation matrix not shown by default, as p = 17 > 12.
## Use print(x, correlation=TRUE)  or
##     vcov(x)        if you need it

## convergence code: 0
```

```
## Model failed to converge with max|grad| = 0.0103859 (tol = 0.001, component 1)
```

##### 1.3. Genome size effect on DBD or EC

Model with the interaction between genus genome size and non focal genera diversity as the predictor for focal lineage diversity (fixed effect)
Model 1 = glmer(nb__ASV~nb__genus*size+(nb__genus|genus__type)+(1|sample))
Model 2 (model with environment as a random effect) :
Model 2 = glmer(nb__ASV~nb__genus*size+(nb__genus|genus__type/empo__3)+(nb__genus|empo__3)+(1|sample))

Genome size effect is not significant in both models.

###### 1.3.1. Model 1.

```
## Generalized linear mixed model fit by maximum likelihood (Laplace
##   Approximation) [glmerMod]
##  Family: poisson  ( log )
## Formula: nb_ASV ~ nb_genus + (1 | genus_type)
##    Data: datsc_size
## Control: glmerControl(optimizer = "bobyqa")
##
##      AIC      BIC   logLik deviance df.resid
##  2368697  2368731 -1184345  2368691   712683
##
## Scaled residuals:
##     Min      1Q  Median      3Q     Max
## -3.1185 -0.4458 -0.1544  0.4320  5.5805
##
## Random effects:
##  Groups     Name        Variance Std.Dev.
##  genus_type (Intercept) 0.5072   0.7122
## Number of obs: 712686, groups:  genus_type, 576
##
## Fixed effects:
##               Estimate Std. Error z value Pr(>|z|)
## (Intercept)  0.6745515  0.0291069  23.175   <2e-16 ***
## nb_genus    -0.0054999  0.0005967  -9.217   <2e-16 ***
## ---
## Signif. codes:  0 '***' 0.001 '**' 0.01 '*' 0.05 '.' 0.1 ' ' 1
## convergence code: 0
## Model is nearly unidentifiable: very large eigenvalue
##  - Rescale variables?
```

###### 1.3.2. Model 2

```
## Generalized linear mixed model fit by maximum likelihood (Laplace
##   Approximation) [glmerMod]
##  Family: poisson  ( log )
## Formula: nb_ASV ~ nb_genus + (nb_genus | genus_type) + (1 | empo_3)
##    Data: datsc_size
## Control: glmerControl(optimizer = "bobyqa")
```

```
##
##      AIC      BIC   logLik deviance df.resid
##  2333229  2333298 -1166608  2333217   704357
##
## Scaled residuals:
##     Min      1Q  Median      3Q     Max
## -3.2877 -0.4451 -0.1510  0.4172  5.1661
##
## Random effects:
##  Groups     Name        Variance  Std.Dev. Corr
##  genus_type (Intercept) 0.4969558 0.70495
##             nb_genus    0.0005646 0.02376  0.84
##  empo_3     (Intercept) 0.0114327 0.10692
## Number of obs: 704363, groups:  genus_type, 576; empo_3, 17
##
## Fixed effects:
##              Estimate Std. Error z value Pr(>|z|)
## (Intercept)  0.661654   0.039154   16.90   <2e-16 ***
## nb_genus    -0.042025   0.001943  -21.63   <2e-16 ***
## ---
## Signif. codes:  0 '***' 0.001 '**' 0.01 '*' 0.05 '.' 0.1 ' ' 1
## convergence code: 0
## Model failed to converge with max|grad| = 0.0176292 (tol = 0.002, component 1)
## Model is nearly unidentifiable: very large eigenvalue
##  - Rescale variables?
```

### 2. Poisson distribution data (model 3)

#### 2.1. Basic GLMM

Models with focal lineage diversity as a function of community diversity in genus:ASV level, with genus name (genus_type) and EMP sample id (sample) as random effects in model 3 :
model = glmer(nb_ASV~nb_genus+(nb_genus|genus_type)+(1|sample))

##### 2.1.1. Model 3

Significative positive effect of community diversity on focal lineage diversity

```
## Generalized linear mixed model fit by maximum likelihood (Laplace
##   Approximation) [glmerMod]
##  Family: poisson  ( log )
## Formula: nb_ASV ~ nb_genus + (nb_genus - 1 | genus_type)
##    Data: datsc1
## Control: glmerControl(optimizer = "bobyqa", optCtrl = list(maxfun = 1e+05))
##
##       AIC       BIC    logLik  deviance  df.resid
##  826145.7  826177.4 -413069.9  826139.7    287677
##
## Scaled residuals:
##     Min      1Q  Median      3Q     Max
## -0.7992 -0.4273 -0.4195  0.3619 13.6922
```

```
##
## Random effects:
##  Groups     Name     Variance Std.Dev.
##  genus_type nb_genus 0.000462 0.02149
## Number of obs: 287680, groups:  genus_type, 1128
##
## Fixed effects:
##              Estimate Std. Error z value Pr(>|z|)
## (Intercept)  0.420660   0.001512 278.201  < 2e-16 ***
## nb_genus    -0.011984   0.001829  -6.552 5.69e-11 ***
## ---
## Signif. codes:  0 '***' 0.001 '**' 0.01 '*' 0.05 '.' 0.1 ' ' 1
```

###### 2.1.2. Model 3 + DBD

Significative positive effect of community diversity on focal lineage diversity

```
## Generalized linear mixed model fit by maximum likelihood (Laplace
##   Approximation) [glmerMod]
##  Family: poisson  ( log )
## Formula: nb_ASV ~ nb_genus + (nb_genus | genus_type) + (1 | sample)
##    Data: datsc1
## Control: glmerControl(optimizer = "bobyqa", optCtrl = list(maxfun = 2e+05))
##
##       AIC       BIC    logLik  deviance  df.resid
## 1565861.8 1565929.2 -782924.9 1565849.8    565360
##
## Scaled residuals:
##     Min      1Q  Median      3Q     Max
## -3.4035 -0.3289 -0.1094  0.2103  6.0892
##
## Random effects:
##  Groups     Name        Variance  Std.Dev. Corr
##  sample     (Intercept) 2.609e-04 0.016152
##  genus_type (Intercept) 1.302e-01 0.360837
##             nb_genus    6.674e-05 0.008169 0.91
## Number of obs: 565366, groups:  sample, 2000; genus_type, 1128
##
## Fixed effects:
##             Estimate Std. Error z value Pr(>|z|)
## (Intercept) 0.197417   0.011000   17.95   <2e-16 ***
## nb_genus    0.016332   0.001423   11.48   <2e-16 ***
## ---
## Signif. codes:  0 '***' 0.001 '**' 0.01 '*' 0.05 '.' 0.1 ' ' 1
## convergence code: 0
## Model failed to converge with max|grad| = 0.00354163 (tol = 0.001, component 1)
## Model is nearly unidentifiable: very large eigenvalue
##  - Rescale variables?
```

###### 2.1.3. Model 3 + EC

Significative negative effect of community diversity on focal lineage diversity

```
##
```

```
## Call:
## lm(formula = nb_ASV ~ nb_genus, data = datsc1)
##
## Residuals:
##     Min      1Q  Median      3Q     Max
## -0.2664 -0.2400 -0.2306 -0.2197 10.7818
##
## Coefficients:
##              Estimate Std. Error t value Pr(>|t|)
## (Intercept)  1.233847   0.001762  700.22  < 2e-16 ***
## nb_genus    -0.010820   0.001762   -6.14 8.26e-10 ***
## ---
## Signif. codes:  0 '***' 0.001 '**' 0.01 '*' 0.05 '.' 0.1 ' ' 1
##
## Residual standard error: 0.6362 on 130374 degrees of freedom
## Multiple R-squared:  0.0002891,  Adjusted R-squared:  0.0002814
## F-statistic:  37.7 on 1 and 130374 DF,  p-value: 8.259e-10
```

#### 2.2. Genome size effect on DBD or EC

Model with the interaction between genus genome size and non focal genera diversity as a predictor to focal lineage diversity (fixed effect)
Model = glmer(nb__ASV~nb__genus*size+(nb__genus|genus__type)+(1|sample))

The interaction between genome size and community diversity does not improve the fit (LRT between the full model and the model without the interaction, p=0.01643, dAIC=4)

```
## Generalized linear mixed model fit by maximum likelihood (Laplace
##   Approximation) [glmerMod]
##  Family: poisson  ( log )
## Formula: nb_ASV ~ nb_genus + size + (nb_genus - 1 | genus_type)
##    Data: datsc_size
## Control: glmerControl(optimizer = "bobyqa")
##
##       AIC       BIC    logLik  deviance  df.resid
##  637249.5  637290.7 -318620.8  637241.5    219254
##
## Scaled residuals:
##     Min      1Q  Median      3Q     Max
## -1.1822 -0.4549 -0.3509  0.2114 12.5846
##
## Random effects:
##  Groups     Name     Variance  Std.Dev.
##  genus_type nb_genus 0.0003651 0.01911
## Number of obs: 219258, groups:  genus_type, 576
##
## Fixed effects:
##              Estimate Std. Error z value Pr(>|z|)
## (Intercept)  0.440177   0.001722 255.594  < 2e-16 ***
## nb_genus    -0.011842   0.002049  -5.781 7.44e-09 ***
## size         0.135083   0.001579  85.528  < 2e-16 ***
## ---
## Signif. codes:  0 '***' 0.001 '**' 0.01 '*' 0.05 '.' 0.1 ' ' 1
```
