## Supplementary material for "Does diversity beget diversity in microbiomes?": File S4

### GLMMs output - Delgado dataset

Naima Madi

2019-07-31

#### 1. ASV:Genus

##### 1.1. Model with abiotic variables

```
## Generalized linear mixed model fit by maximum likelihood (Laplace
##   Approximation) [glmerMod]
##  Family: poisson  ( log )
## Formula: nb_ASV ~ Latitude + UV_Light + MDR + NPP2003_2015 + I(Latitude^2) +
##     I(Clay_silt^2) + I(Soil_N^2) + I(UV_Light^2) + I(Soil_C_N_ratio^2) +
##     I(Aridity_Index^2) + I(PSEA^2) + I(MDR^2) + I(NPP2003_2015^2) +
##     (1 | genus_type)
##    Data: datsc
## Control: glmerControl(optimizer = "bobyqa")
##
##      AIC      BIC   logLik deviance df.resid
##  69512.5  69628.4 -34741.3  69482.5    16774
##
## Scaled residuals:
##     Min      1Q  Median      3Q     Max
## -4.4614 -0.5549 -0.1745  0.4129 13.8404
##
## Random effects:
##  Groups     Name        Variance Std.Dev.
##  genus_type (Intercept) 0.4181   0.6466
## Number of obs: 16789, groups:  genus_type, 297
##
## Fixed effects:
##                       Estimate Std. Error z value Pr(>|z|)
## (Intercept)          0.8106803  0.0478975  16.925  < 2e-16 ***
## Latitude            -0.2936950  0.0252425 -11.635  < 2e-16 ***
## UV_Light            -0.1766322  0.0163646 -10.794  < 2e-16 ***
## MDR                  0.0280697  0.0062517   4.490 7.12e-06 ***
## NPP2003_2015        -0.0660638  0.0054224 -12.184  < 2e-16 ***
## I(Latitude^2)       -0.3002591  0.0286991 -10.462  < 2e-16 ***
## I(Clay_silt^2)      -0.0124015  0.0041383  -2.997 0.002728 **
## I(Soil_N^2)         -0.0068009  0.0014196  -4.791 1.66e-06 ***
## I(UV_Light^2)        0.0003397  0.0044216   0.077 0.938755
## I(Soil_C_N_ratio^2)  0.0030466  0.0010596   2.875 0.004037 **
## I(Aridity_Index^2)  -0.0009996  0.0023075  -0.433 0.664866
## I(PSEA^2)            0.0106587  0.0023315   4.572 4.84e-06 ***
## I(MDR^2)             0.0174667  0.0031299   5.581 2.40e-08 ***
```

```
## I(NPP2003_2015^2)   -0.0160078  0.0041349  -3.871 0.000108 ***
## ---
## Signif. codes:  0 '***' 0.001 '**' 0.01 '*' 0.05 '.' 0.1 ' ' 1
## convergence code: 0
## Model is nearly unidentifiable: very large eigenvalue
##  - Rescale variables?
```

##### 1.2. Model with the three first PCs (ENV1, ENV2, ENV3)

```
## Generalized linear mixed model fit by maximum likelihood (Laplace
##   Approximation) [glmerMod]
##  Family: poisson  ( log )
## Formula: nb_ASV ~ nb_genus + ENV1 + ENV2 + (nb_genus | genus_type) + (1 |
##     sample)
##    Data: datsc1
## Control: glmerControl(optimizer = "bobyqa")
##
##      AIC      BIC   logLik deviance df.resid
##  67680.7  67742.5 -33832.4  67664.7    16781
##
## Scaled residuals:
##     Min      1Q  Median      3Q     Max
## -4.9136 -0.5257 -0.1539  0.4079 13.0634
##
## Random effects:
##  Groups     Name        Variance Std.Dev. Corr
##  genus_type (Intercept) 0.413305 0.64289
##             nb_genus    0.028520 0.16888  0.22
##  sample     (Intercept) 0.005796 0.07613
## Number of obs: 16789, groups:  genus_type, 297; sample, 186
##
## Fixed effects:
##              Estimate Std. Error z value Pr(>|z|)
## (Intercept)  0.487117   0.040650  11.983  < 2e-16 ***
## nb_genus     0.064483   0.016518   3.904 9.47e-05 ***
## ENV1        -0.065343   0.007514  -8.696  < 2e-16 ***
## ENV2        -0.029836   0.006991  -4.268 1.98e-05 ***
## ---
## Signif. codes:  0 '***' 0.001 '**' 0.01 '*' 0.05 '.' 0.1 ' ' 1
```

#### 2. Genus:Family

##### 2.1. Model with abiotic variables

```
## Generalized linear mixed model fit by maximum likelihood (Laplace
##   Approximation) [glmerMod]
##  Family: poisson  ( log )
## Formula: nb_genus ~ nb_family + Latitude + (nb_family | family_type)
##    Data: datsc
## Control: glmerControl(optimizer = "bobyqa")
##
```

```
##      AIC      BIC   logLik deviance df.resid
##  41698.9  41745.1 -20843.4  41686.9    16349
##
## Scaled residuals:
##     Min      1Q  Median      3Q     Max
## -1.7485 -0.1631 -0.0188  0.0935  3.2675
##
## Random effects:
##  Groups      Name        Variance Std.Dev. Corr
##  family_type (Intercept) 0.178438 0.42242
##              nb_family   0.004376 0.06615  0.82
## Number of obs: 16355, groups:  family_type, 187
##
## Fixed effects:
##              Estimate Std. Error z value Pr(>|z|)
## (Intercept)  0.277222   0.034325   8.076 6.67e-16 ***
## nb_family    0.032444   0.009938   3.265   0.0011 **
## Latitude    -0.034974   0.005835  -5.994 2.04e-09 ***
## ---
## Signif. codes:  0 '***' 0.001 '**' 0.01 '*' 0.05 '.' 0.1 ' ' 1
```

##### 2.2. Model with the three first PCs (ENV1, ENV2, ENV3)

```
## Generalized linear mixed model fit by maximum likelihood (Laplace
##   Approximation) [glmerMod]
##  Family: poisson  ( log )
## Formula: nb_genus ~ nb_family + ENV1 + ENV2 + (nb_family | family_type)
##    Data: datsc1
## Control: glmerControl(optimizer = "bobyqa")
##
##      AIC      BIC   logLik deviance df.resid
##  41719.2  41773.1 -20852.6  41705.2    16348
##
## Scaled residuals:
##     Min      1Q  Median      3Q     Max
## -1.7245 -0.1638 -0.0268  0.0951  3.2765
##
## Random effects:
##  Groups      Name        Variance Std.Dev. Corr
##  family_type (Intercept) 0.178848 0.42290
##              nb_family   0.004359 0.06602  0.82
## Number of obs: 16355, groups:  family_type, 187
##
## Fixed effects:
##              Estimate Std. Error z value Pr(>|z|)
## (Intercept)  0.277032   0.034364   8.062 7.53e-16 ***
## nb_family    0.032774   0.010163   3.225  0.00126 **
## ENV1        -0.015649   0.006572  -2.381  0.01725 *
## ENV2         0.019839   0.005970   3.323  0.00089 ***
## ---
## Signif. codes:  0 '***' 0.001 '**' 0.01 '*' 0.05 '.' 0.1 ' ' 1
```

### 3. Family:Order

#### 3.1. Model with abiotic variables

```
## Generalized linear mixed model fit by maximum likelihood (Laplace
##   Approximation) [glmerMod]
##  Family: poisson  ( log )
## Formula: nb_family ~ Latitude + (1 | order_type)
##    Data: datsc
## Control: glmerControl(optimizer = "bobyqa")
##
##      AIC      BIC   logLik deviance df.resid
##  32240.6  32263.1 -16117.3  32234.6    13163
##
## Scaled residuals:
##     Min      1Q  Median      3Q     Max
## -3.1667 -0.0391 -0.0054  0.0095  3.0092
##
## Random effects:
##  Groups     Name        Variance Std.Dev.
##  order_type (Intercept) 0.1953   0.4419
## Number of obs: 13166, groups:  order_type, 160
##
## Fixed effects:
##               Estimate Std. Error z value Pr(>|z|)
## (Intercept)  0.2076332  0.0385928   5.380 7.44e-08 ***
## Latitude    -0.0004861  0.0001899  -2.559   0.0105 *
## ---
## Signif. codes:  0 '***' 0.001 '**' 0.01 '*' 0.05 '.' 0.1 ' ' 1
## convergence code: 0
## Model failed to converge with max|grad| = 0.00981316 (tol = 0.001, component 1)
## Model is nearly unidentifiable: very large eigenvalue
##  - Rescale variables?
```

#### 3.3. Model with the three first PCs (ENV1, ENV2, ENV3)

```
## Generalized linear mixed model fit by maximum likelihood (Laplace
##   Approximation) [glmerMod]
##  Family: poisson  ( log )
## Formula: nb_family ~ nb_order:ENV1 + nb_order:ENV3 + ENV1 + (1 | order_type)
##    Data: datsc1
## Control: glmerControl(optimizer = "bobyqa")
##
##      AIC      BIC   logLik deviance df.resid
##  32141.8  32179.3 -16065.9  32131.8    13161
##
## Scaled residuals:
##      Min       1Q   Median       3Q      Max
## -2.54656 -0.05952 -0.01904  0.04660  2.95171
##
## Random effects:
##  Groups     Name        Variance Std.Dev.
##  order_type (Intercept) 0.1974   0.4442
```

```
## Number of obs: 13166, groups:  order_type, 160
##
## Fixed effects:
##                Estimate Std. Error z value Pr(>|z|)
## (Intercept)    0.211377   0.038592   5.477 4.32e-08 ***
## ENV1          -0.025882   0.007192  -3.598  0.00032 ***
## nb_order:ENV1  0.040433   0.005756   7.025 2.14e-12 ***
## nb_order:ENV3  0.023020   0.004807   4.789 1.68e-06 ***
## ---
## Signif. codes:  0 '***' 0.001 '**' 0.01 '*' 0.05 '.' 0.1 ' ' 1
```

### 4. Order:Class

**No factor (biotic or abiotic) is significant**

### 5. Class:Phylum

#### 5.1. Model with all abiotic variables

```
## Generalized linear mixed model fit by maximum likelihood (Laplace
##   Approximation) [glmerMod]
##  Family: poisson  ( log )
## Formula: nb_clas ~ nb_phyla + pH + (1 | phyla_type)
##    Data: datsc
## Control: glmerControl(optimizer = "bobyqa")
##
##      AIC      BIC   logLik deviance df.resid
##  10509.9  10534.5  -5250.9  10501.9     3469
##
## Scaled residuals:
##      Min       1Q   Median       3Q      Max
## -2.40114 -0.24247 -0.02056  0.19578  2.28775
##
## Random effects:
##  Groups     Name        Variance Std.Dev.
##  phyla_type (Intercept) 0.4592   0.6777
## Number of obs: 3473, groups:  phyla_type, 36
##
## Fixed effects:
##             Estimate Std. Error z value Pr(>|z|)
## (Intercept)  0.49679    0.11956   4.155 3.25e-05 ***
## nb_phyla     0.03152    0.01006   3.132  0.00174 **
## pH           0.07365    0.01017   7.243 4.37e-13 ***
## ---
## Signif. codes:  0 '***' 0.001 '**' 0.01 '*' 0.05 '.' 0.1 ' ' 1
```

#### 5.2. Model with PCs

```
## Generalized linear mixed model fit by maximum likelihood (Laplace
##   Approximation) [glmerMod]
```

```
##  Family: poisson  ( log )
## Formula: nb_clas ~ nb_phyla + ENV1 + ENV2 + (1 | phyla_type)
##    Data: datsc1
## Control: glmerControl(optimizer = "bobyqa")
##
##      AIC      BIC   logLik deviance df.resid
##  10530.2  10560.9  -5260.1  10520.2     3468
##
## Scaled residuals:
##      Min       1Q   Median       3Q      Max
## -2.42028 -0.24084 -0.02345  0.17437  2.34184
##
## Random effects:
##  Groups     Name        Variance Std.Dev.
##  phyla_type (Intercept) 0.4584   0.677
## Number of obs: 3473, groups:  phyla_type, 36
##
## Fixed effects:
##             Estimate Std. Error z value Pr(>|z|)
## (Intercept)  0.49619    0.11944   4.154 3.26e-05 ***
## nb_phyla     0.03153    0.01060   2.975  0.00293 **
## ENV1        -0.05143    0.01010  -5.092 3.54e-07 ***
## ENV2        -0.02847    0.01038  -2.744  0.00606 **
## ---
## Signif. codes:  0 '***' 0.001 '**' 0.01 '*' 0.05 '.' 0.1 ' ' 1
```
