## Supplementary figures and images for "Does diversity beget diversity in microbiomes?"

### Figure 2 Supplement 1

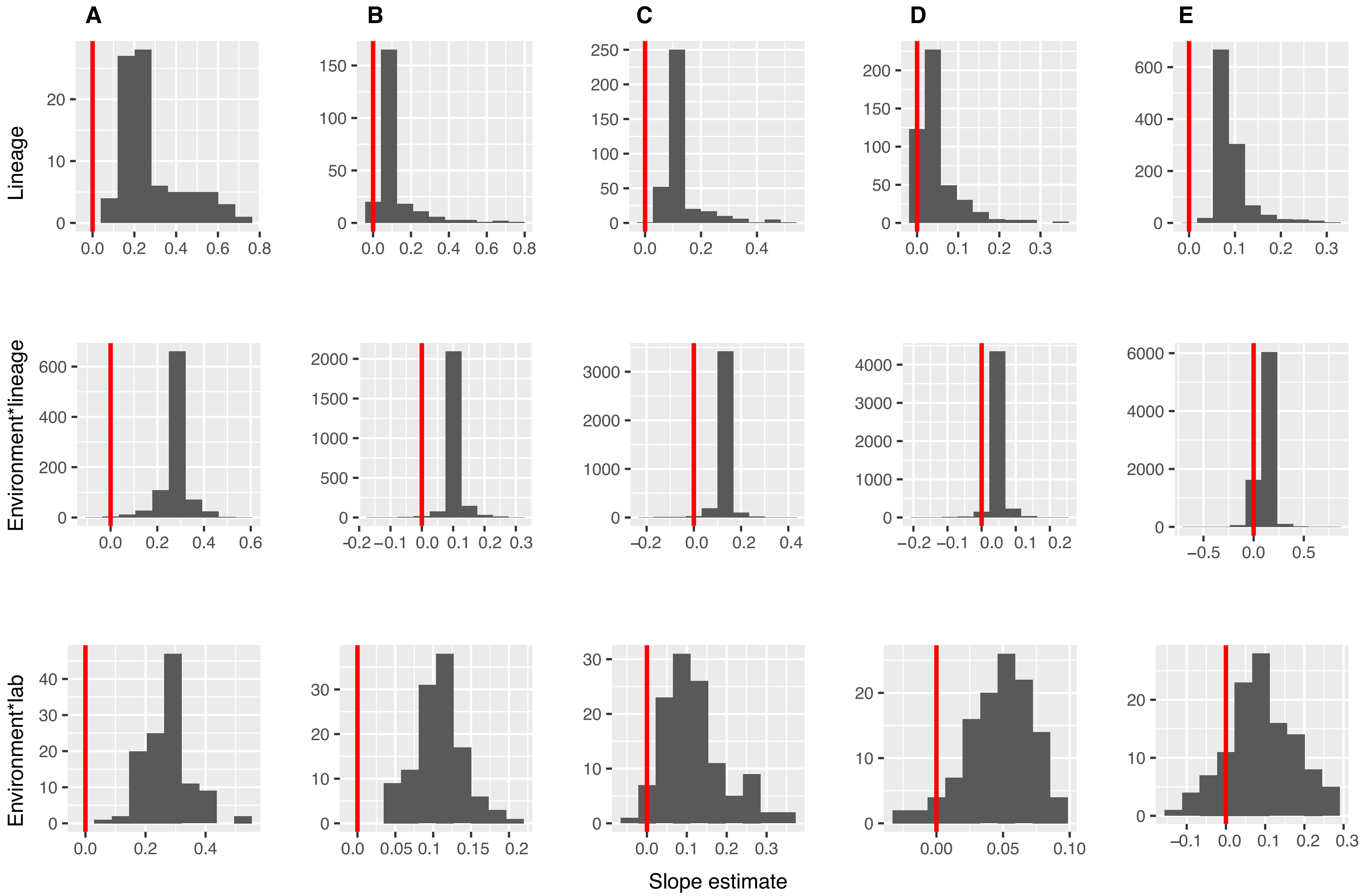

### Figure 2 Supplement 2

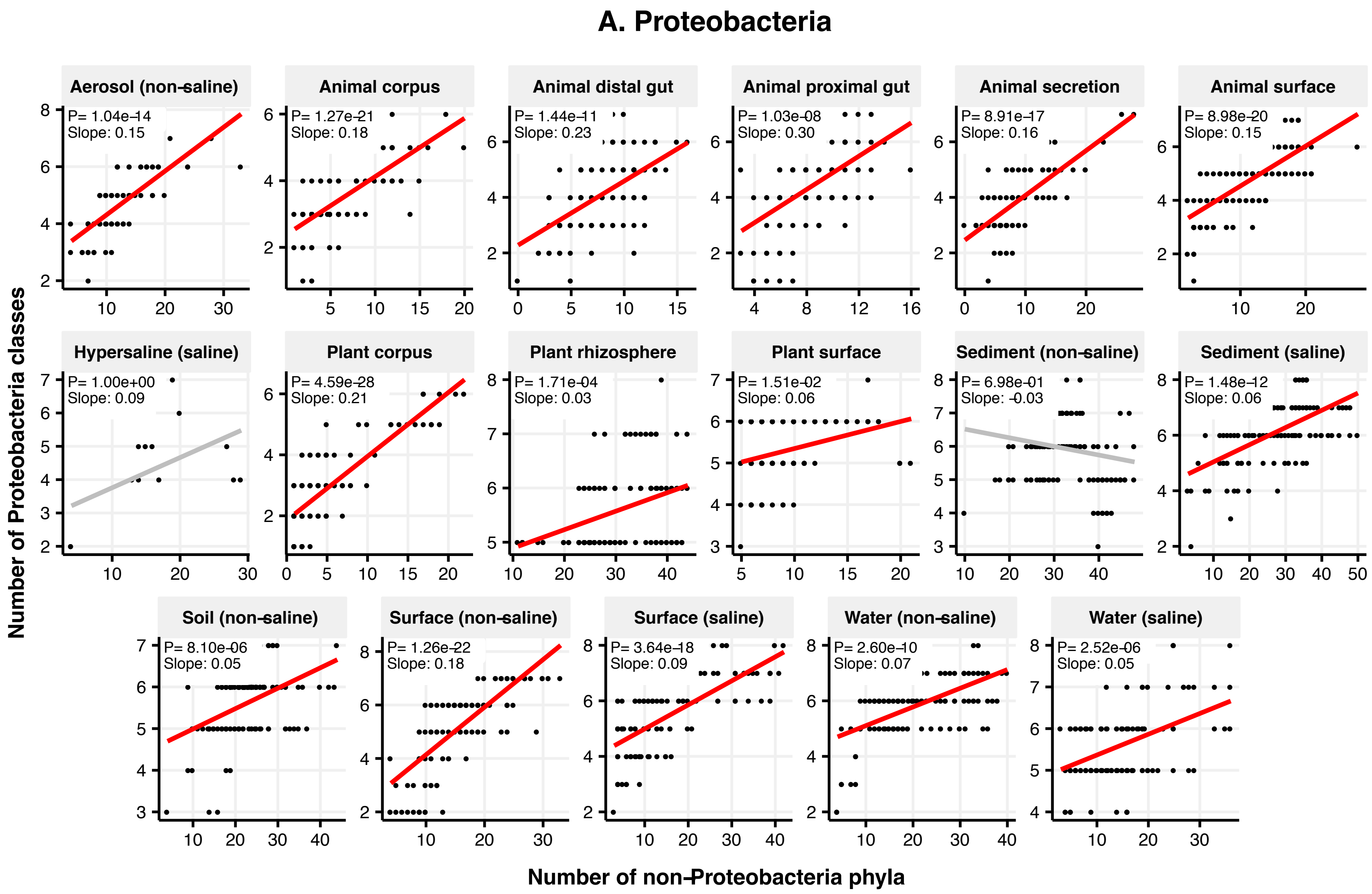

#### B. Bacteroidetes

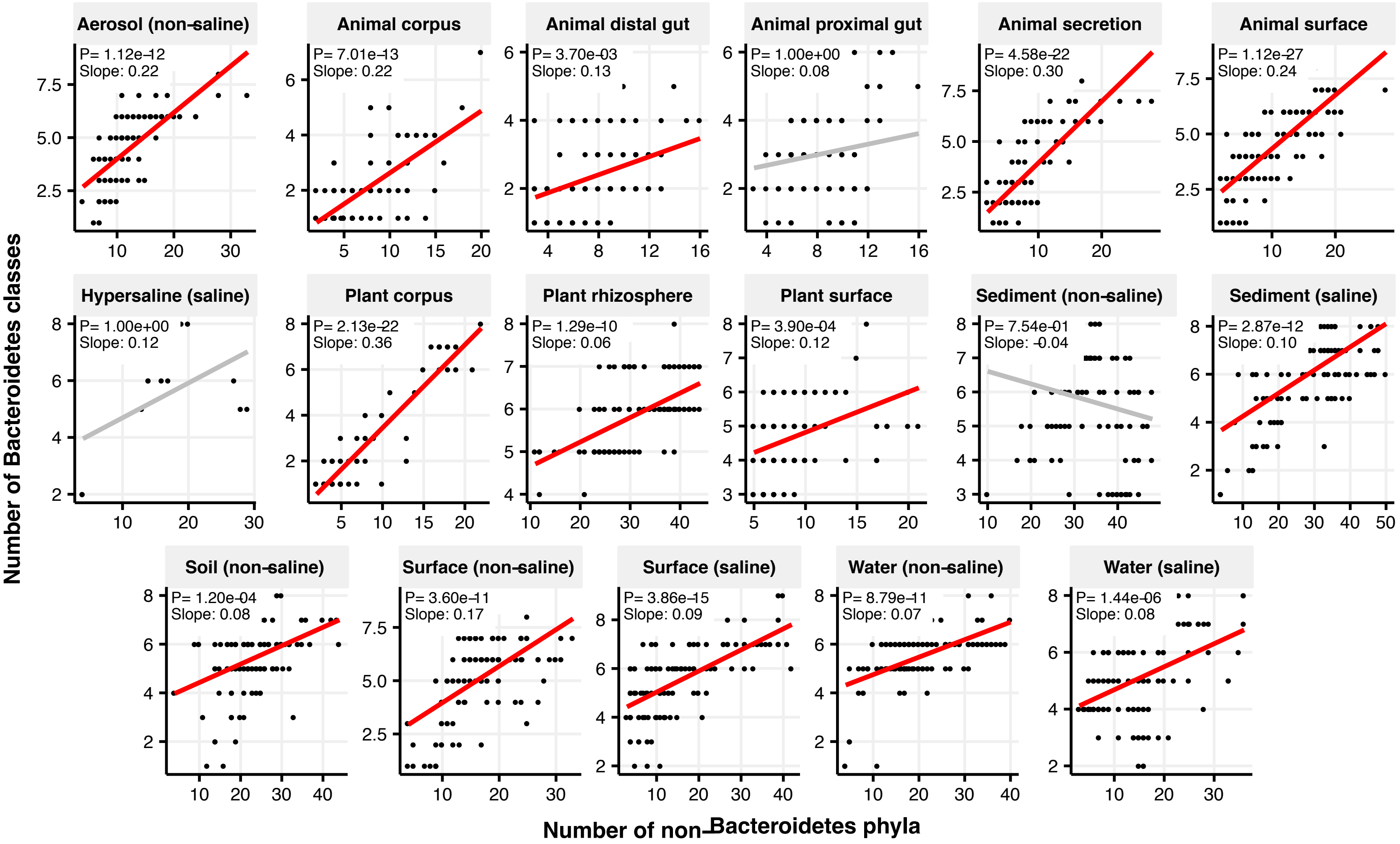

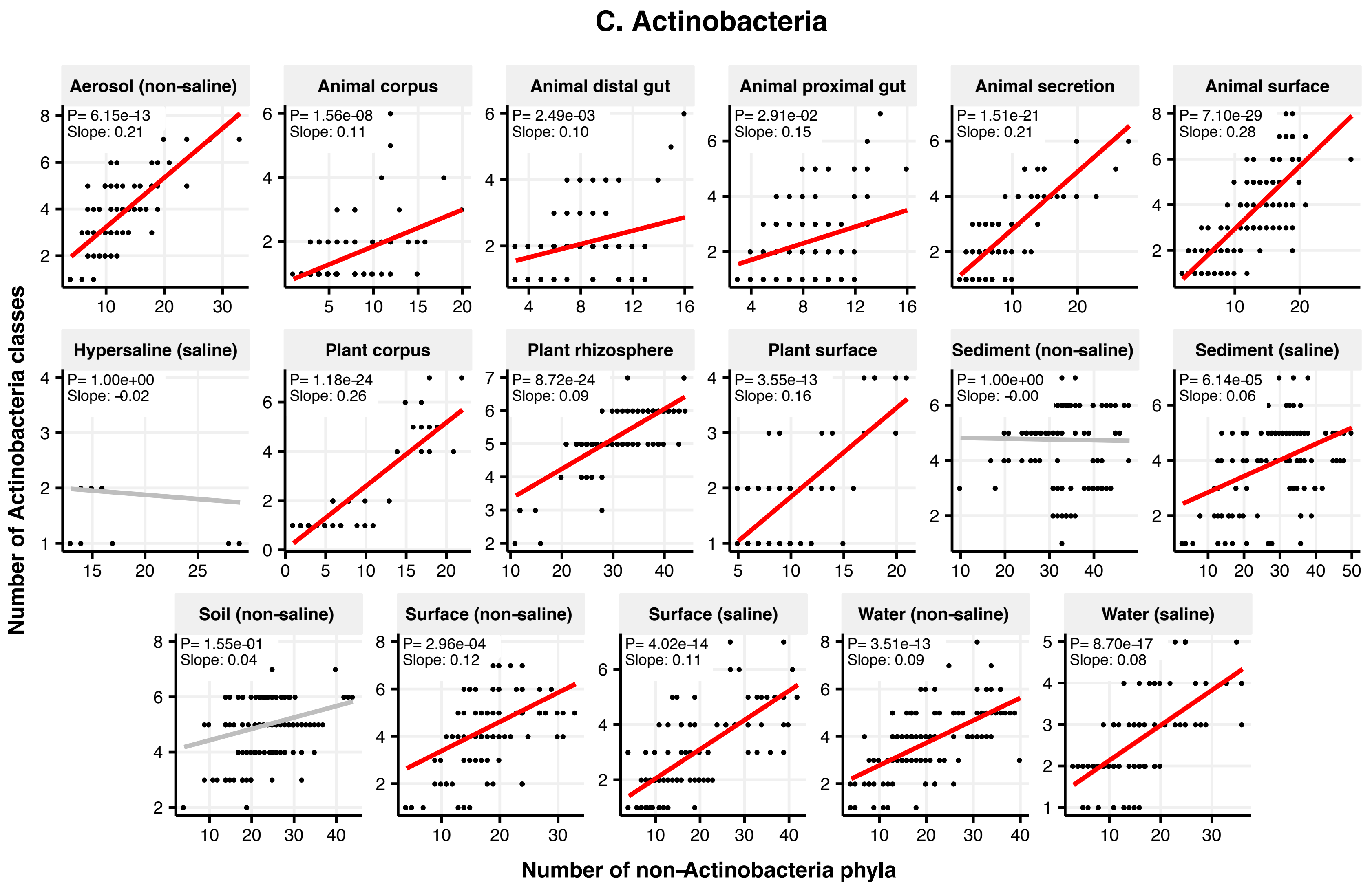

### Figure 2 Supplement 3

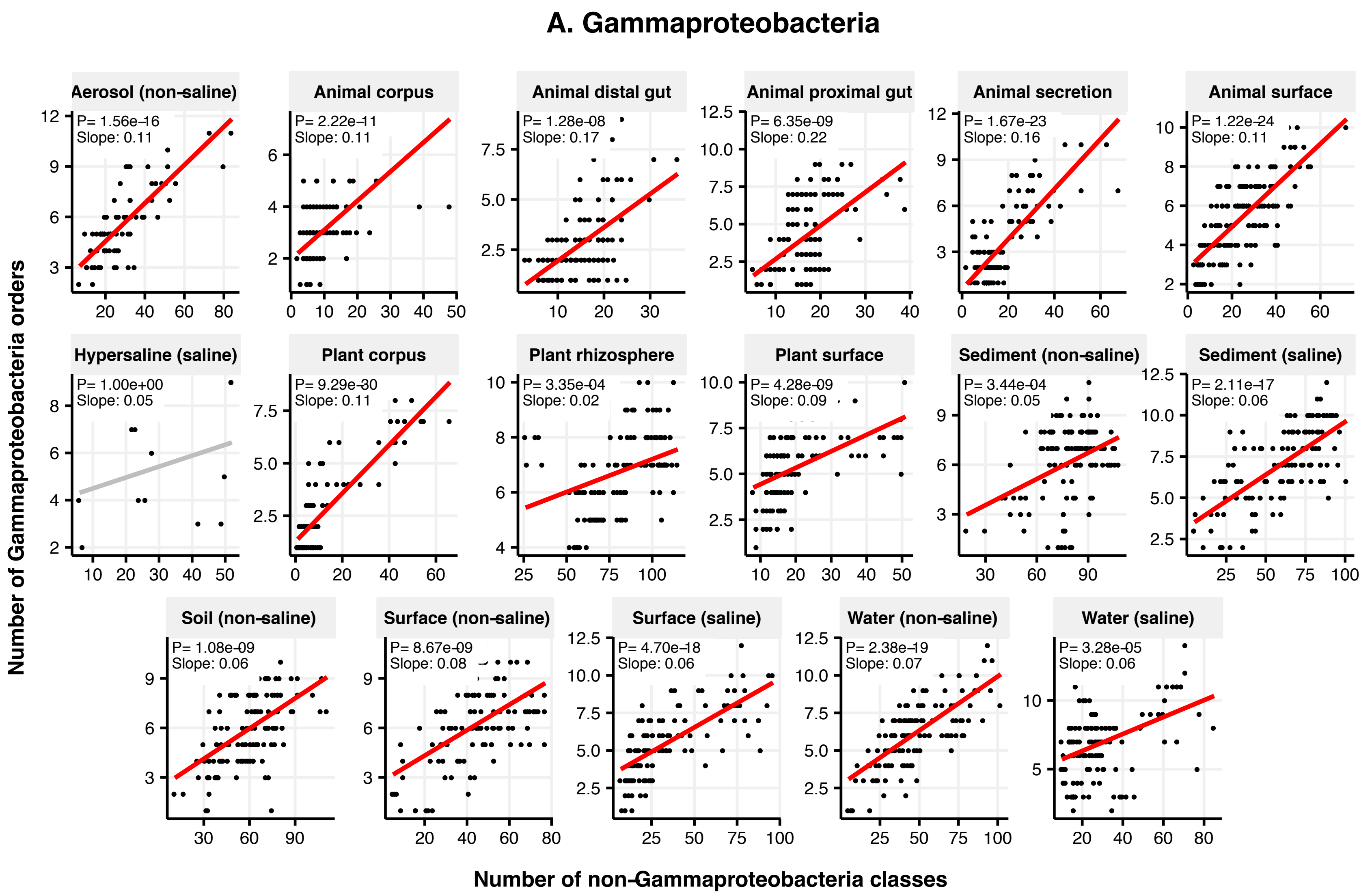

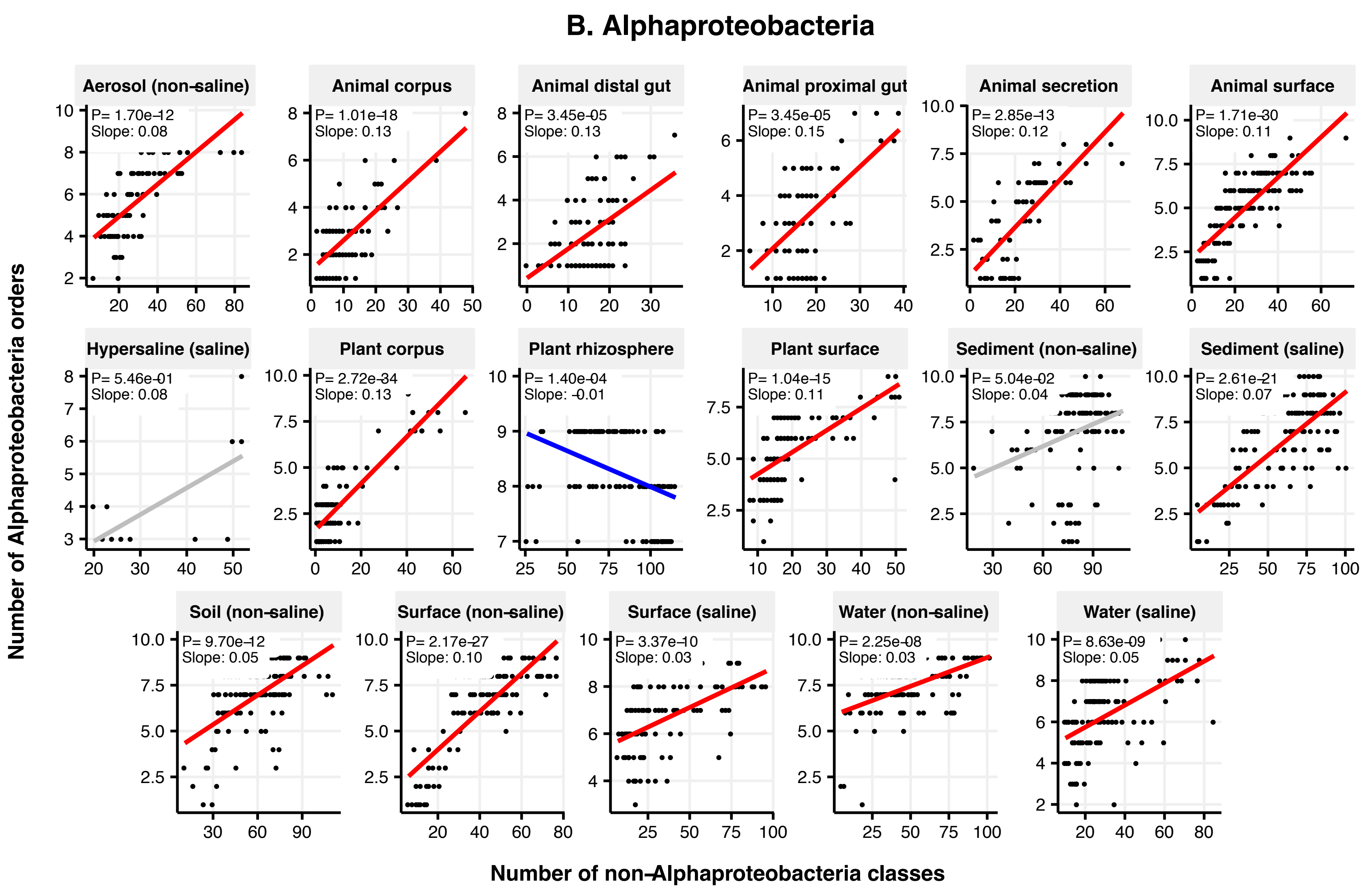

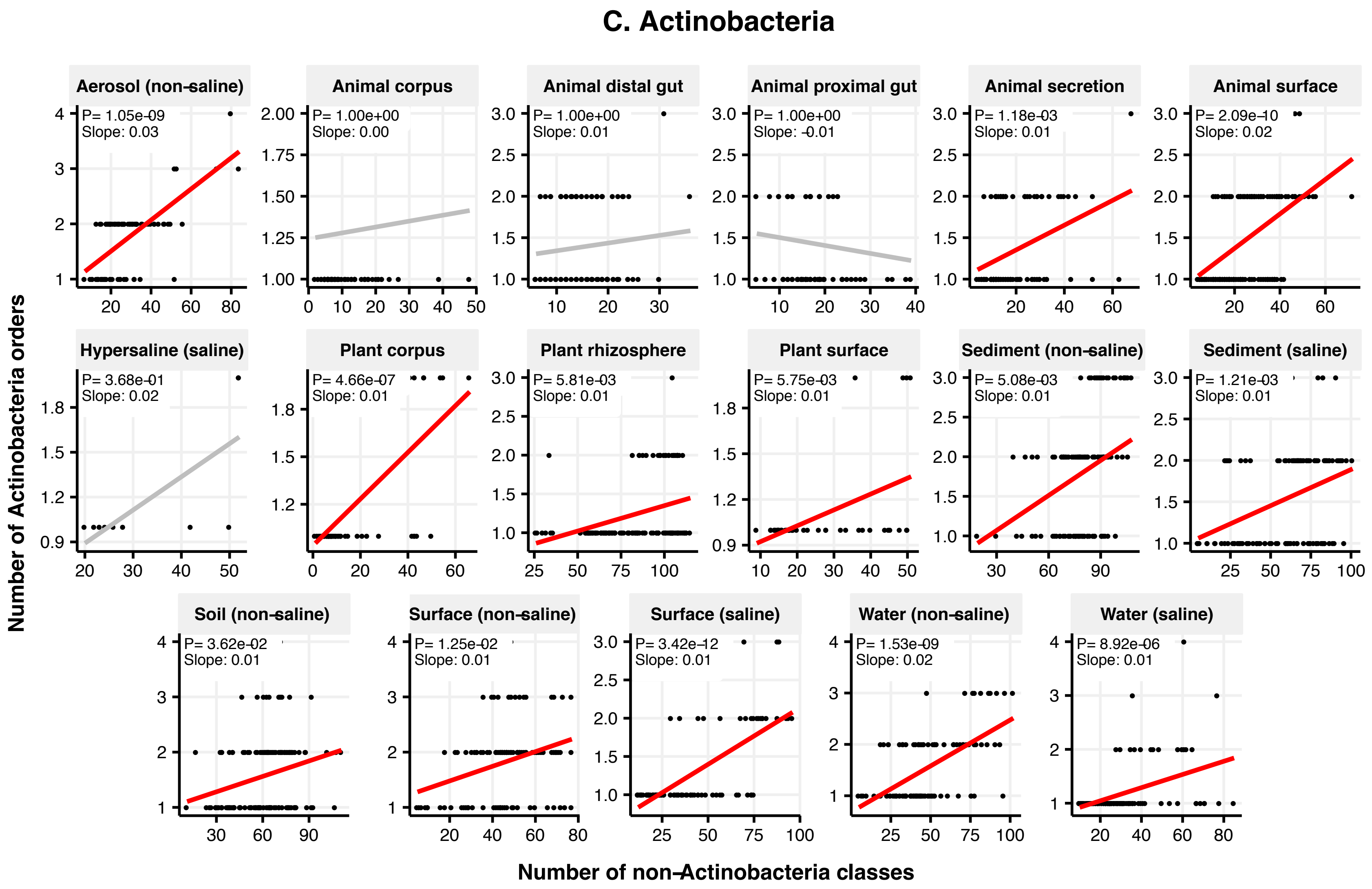

### Figure 2 Supplement 4

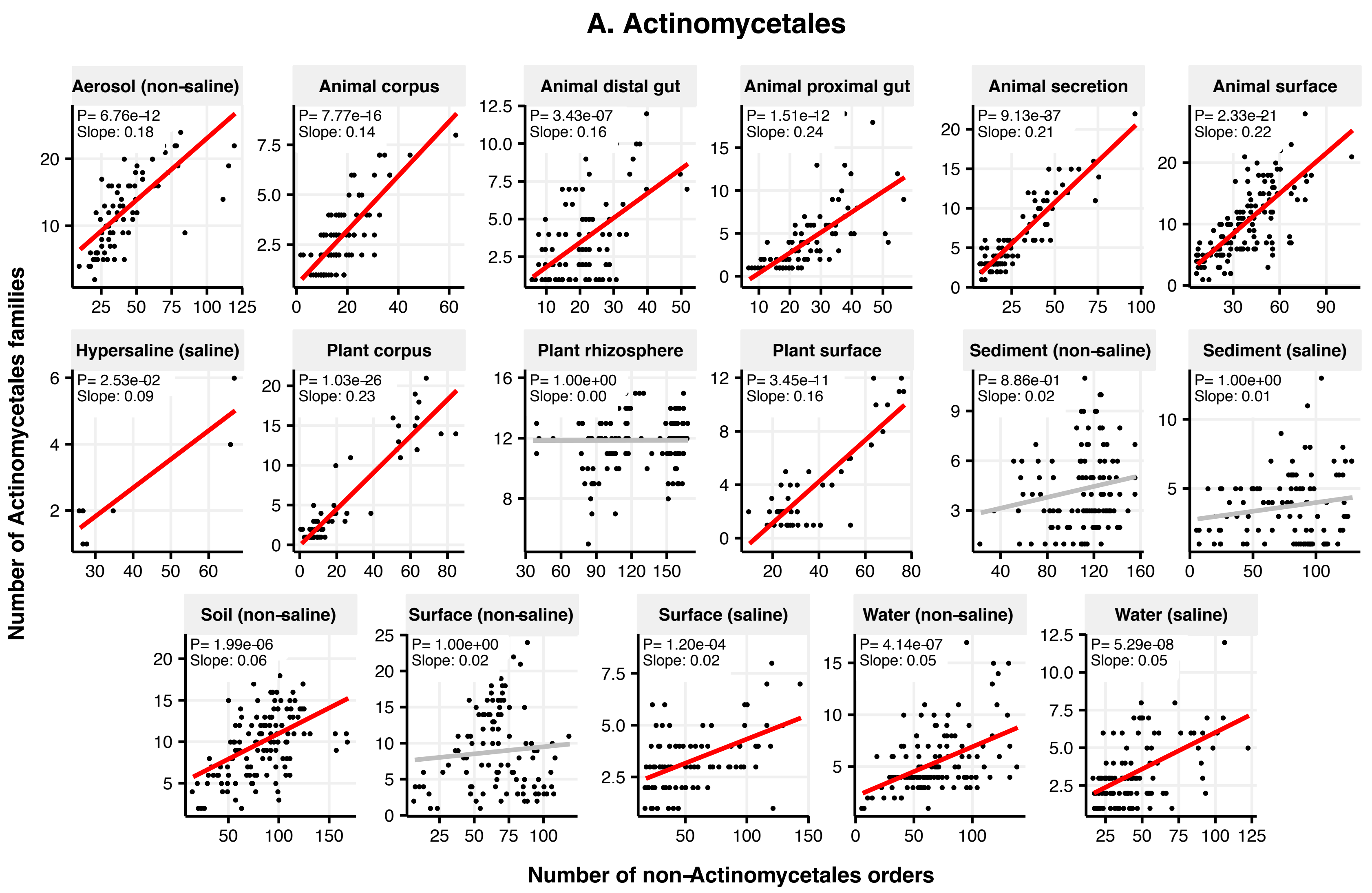

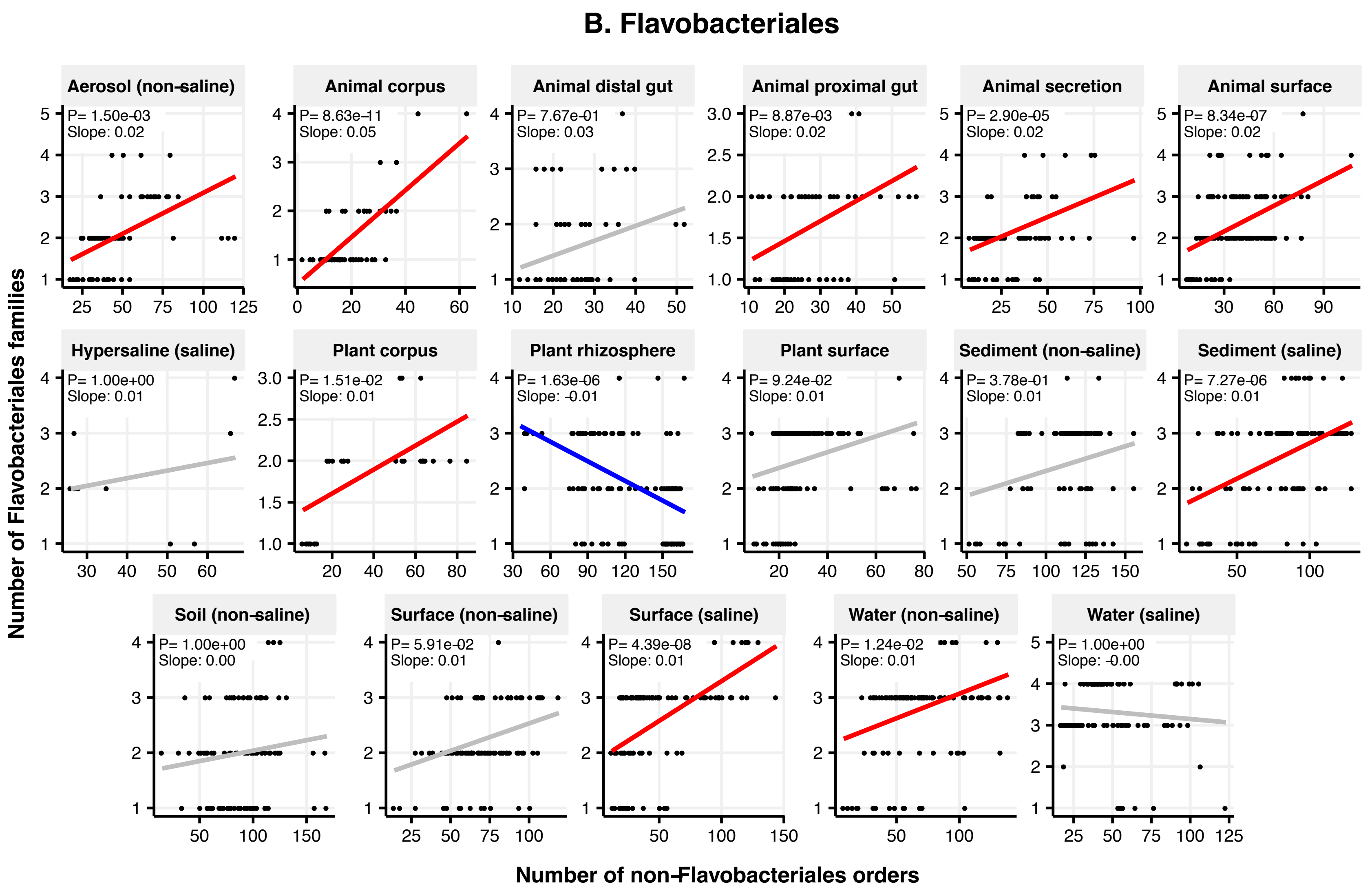

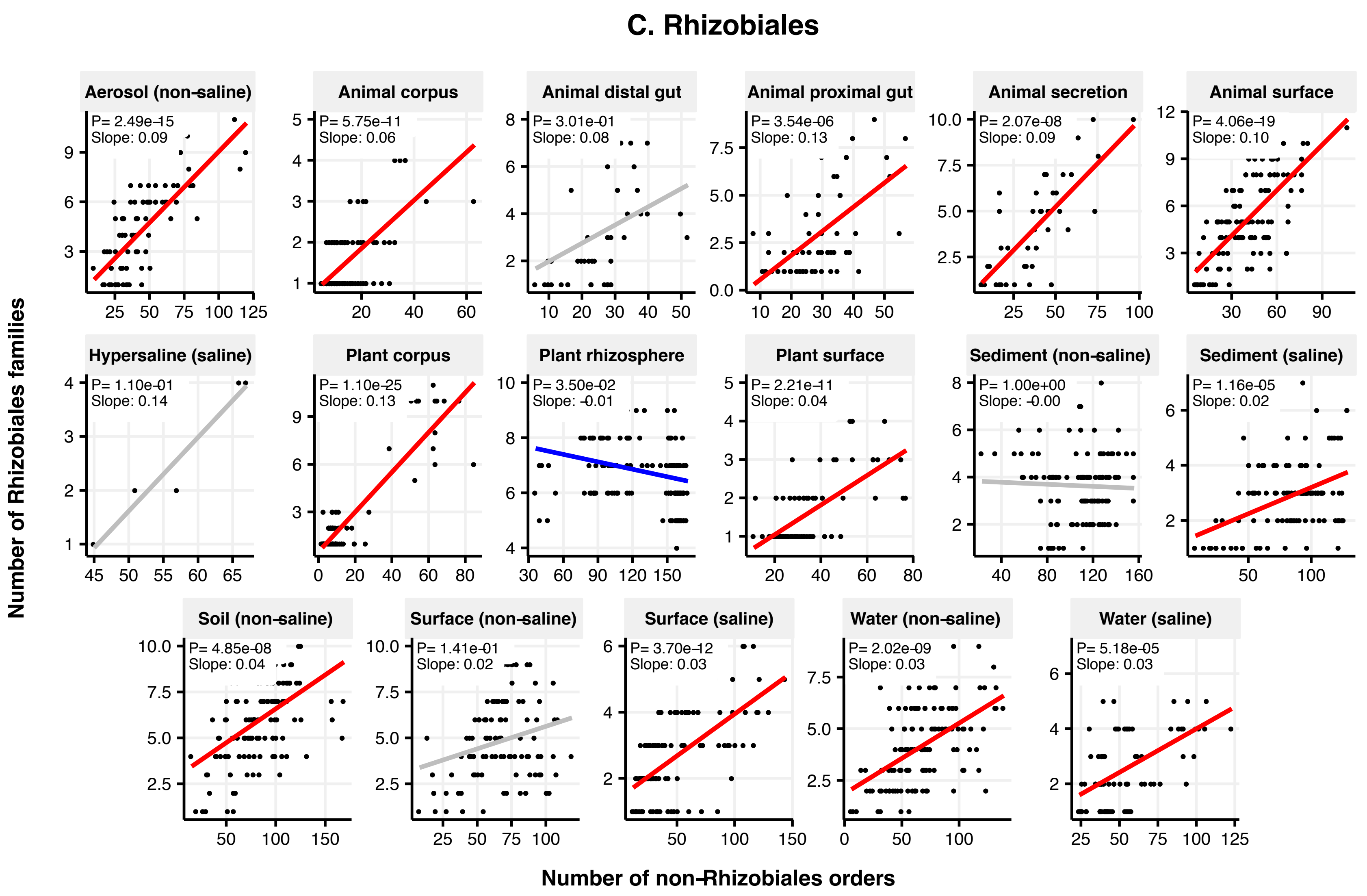

### Figure 2 Supplement 5

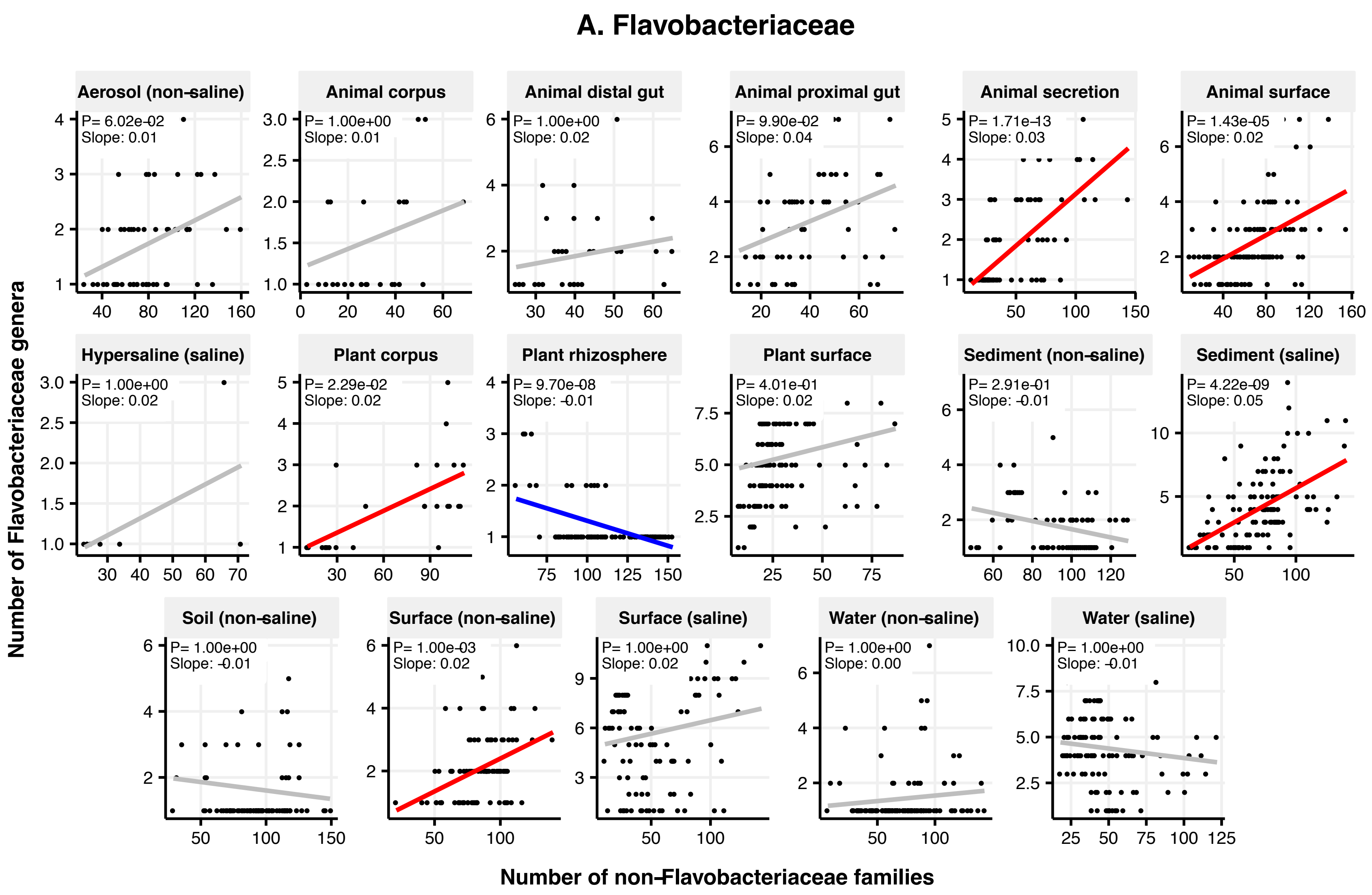

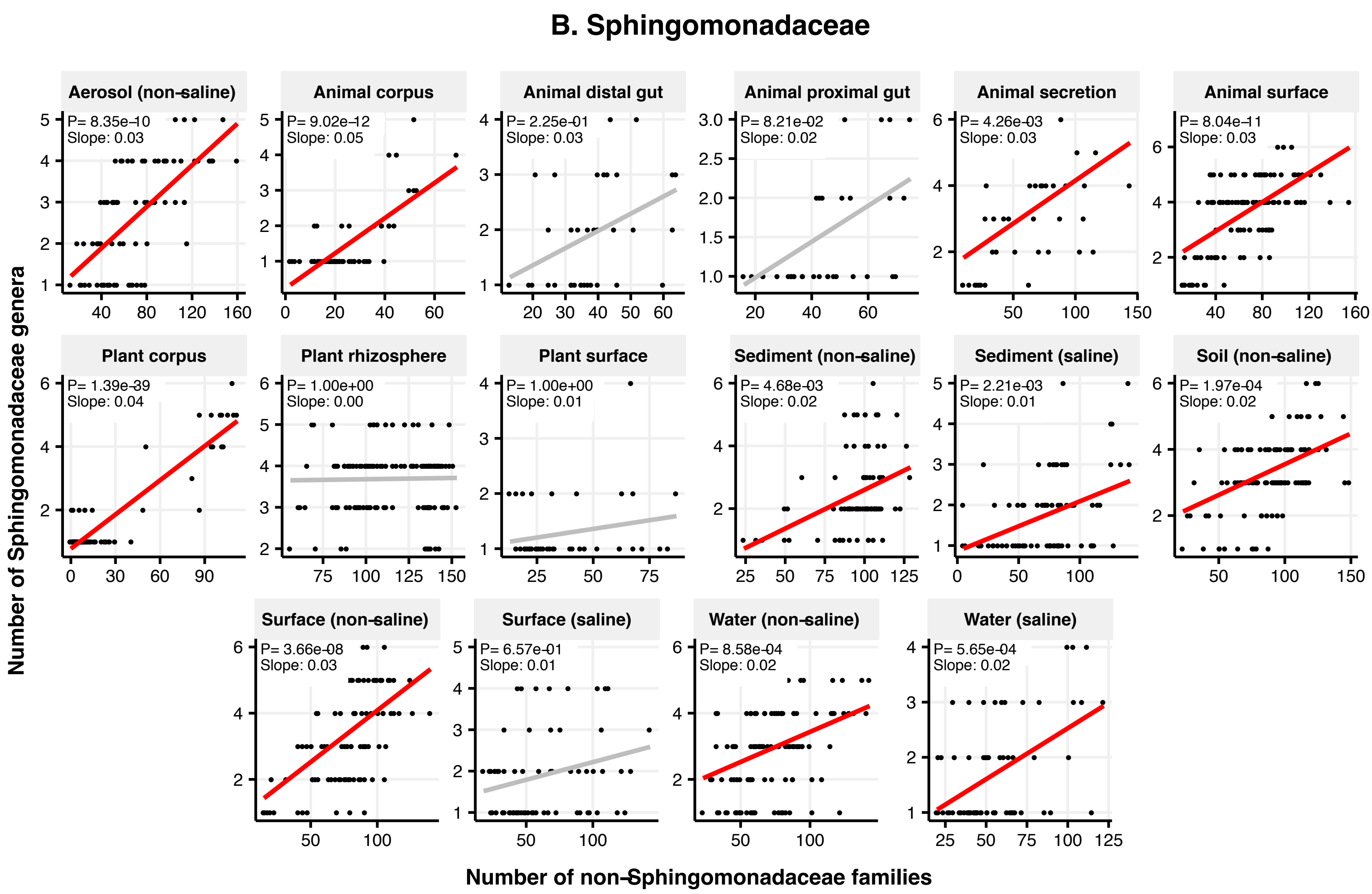

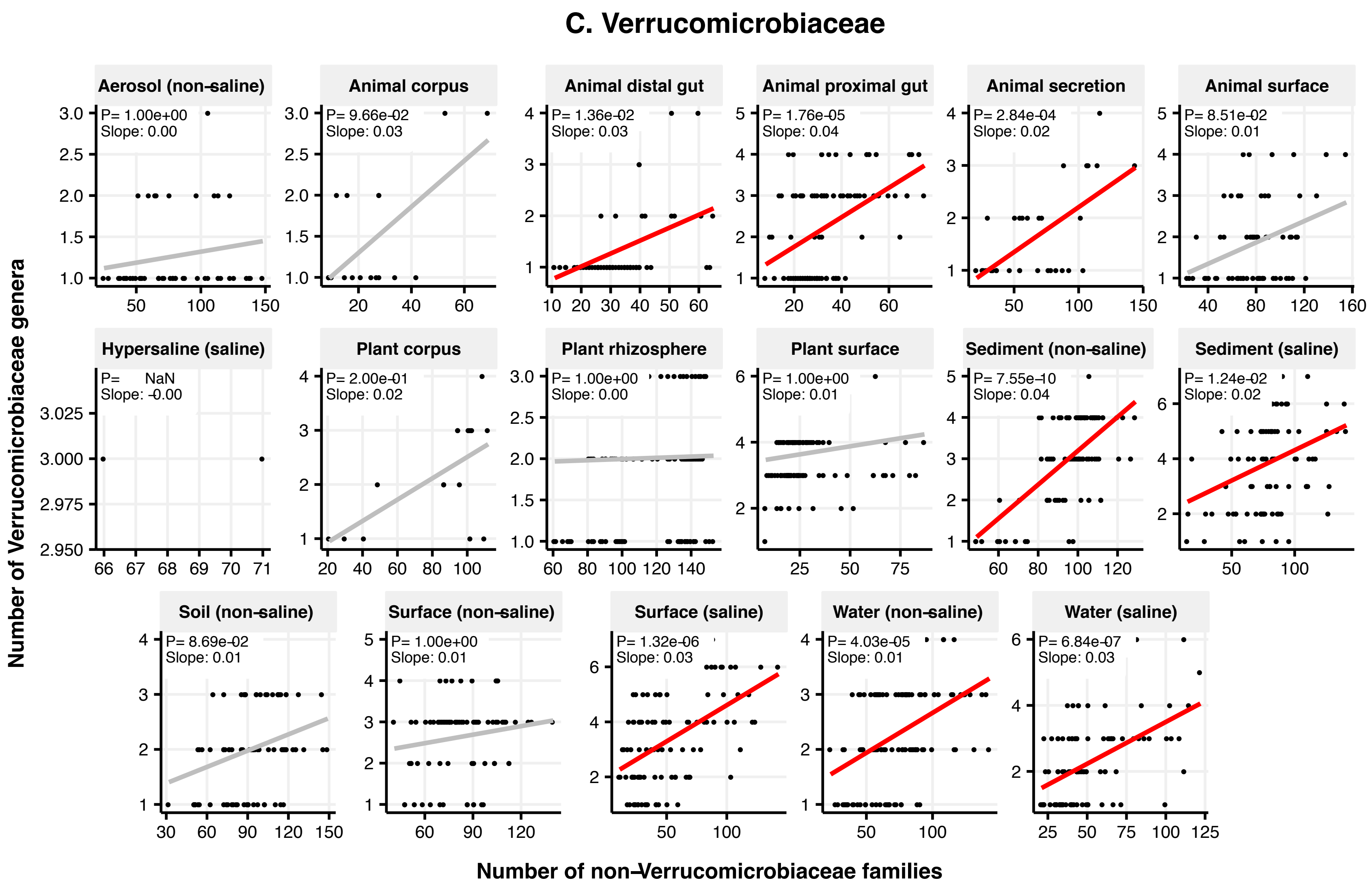

### Figure 2 Supplement 6

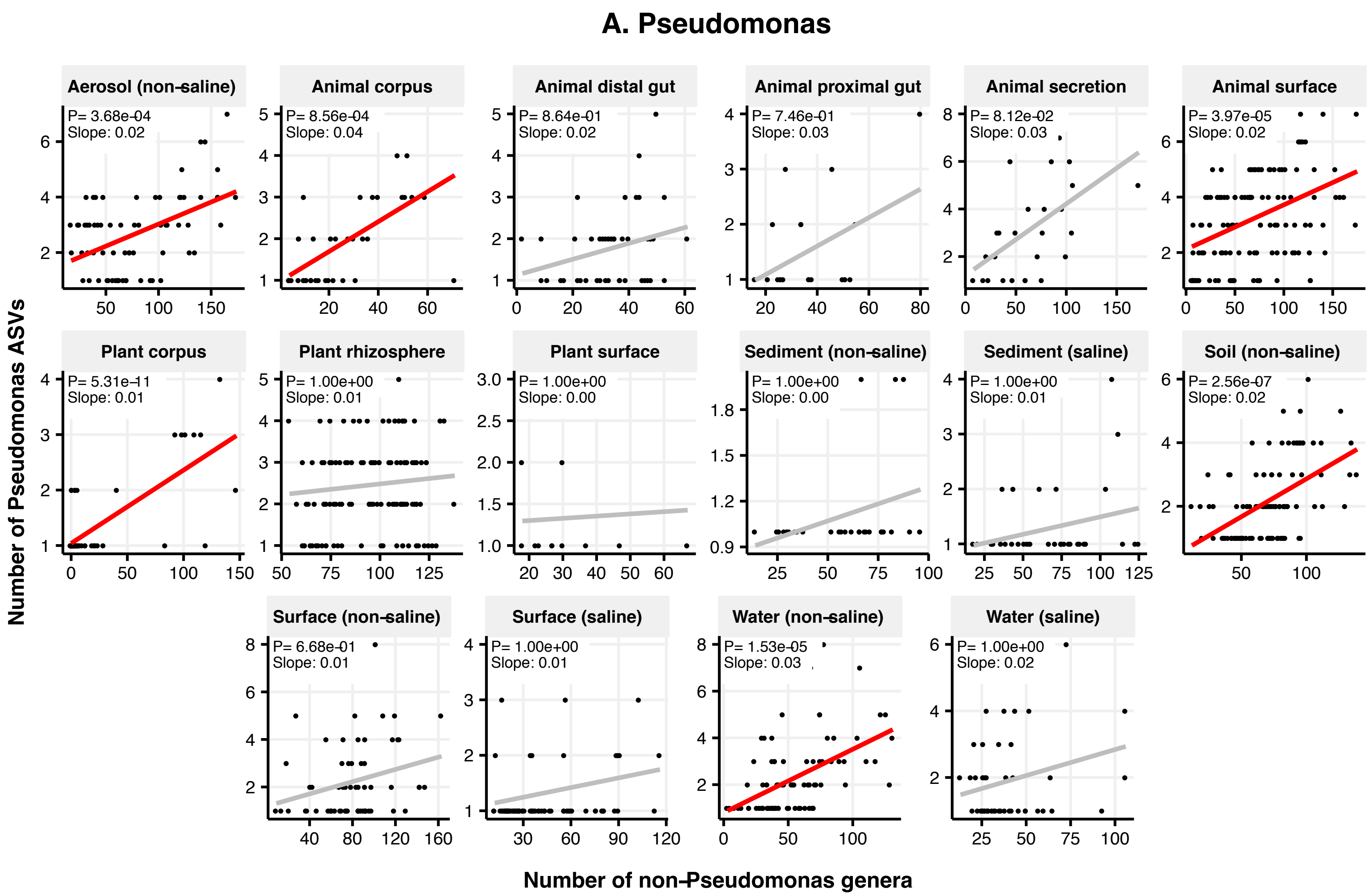

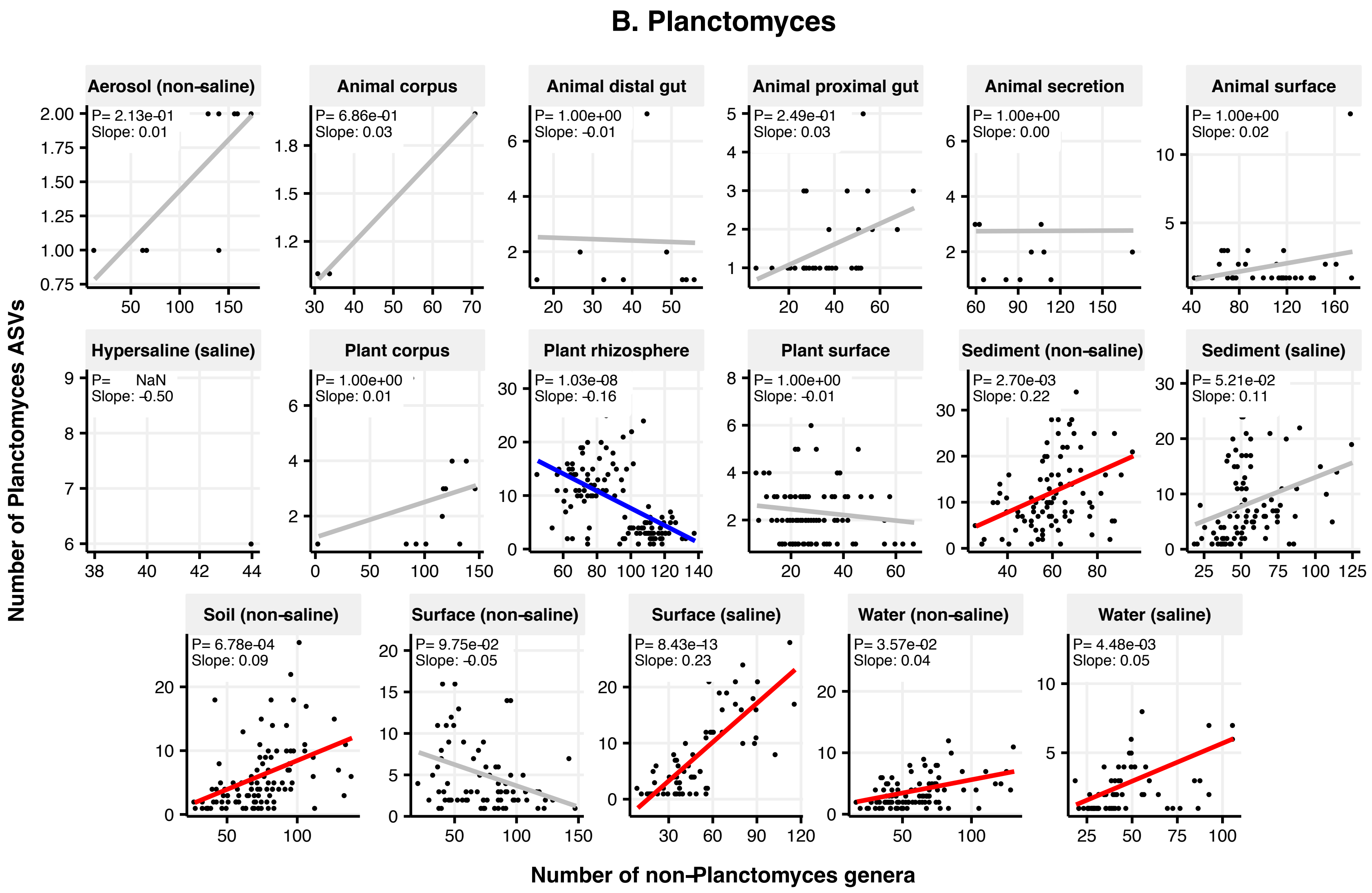

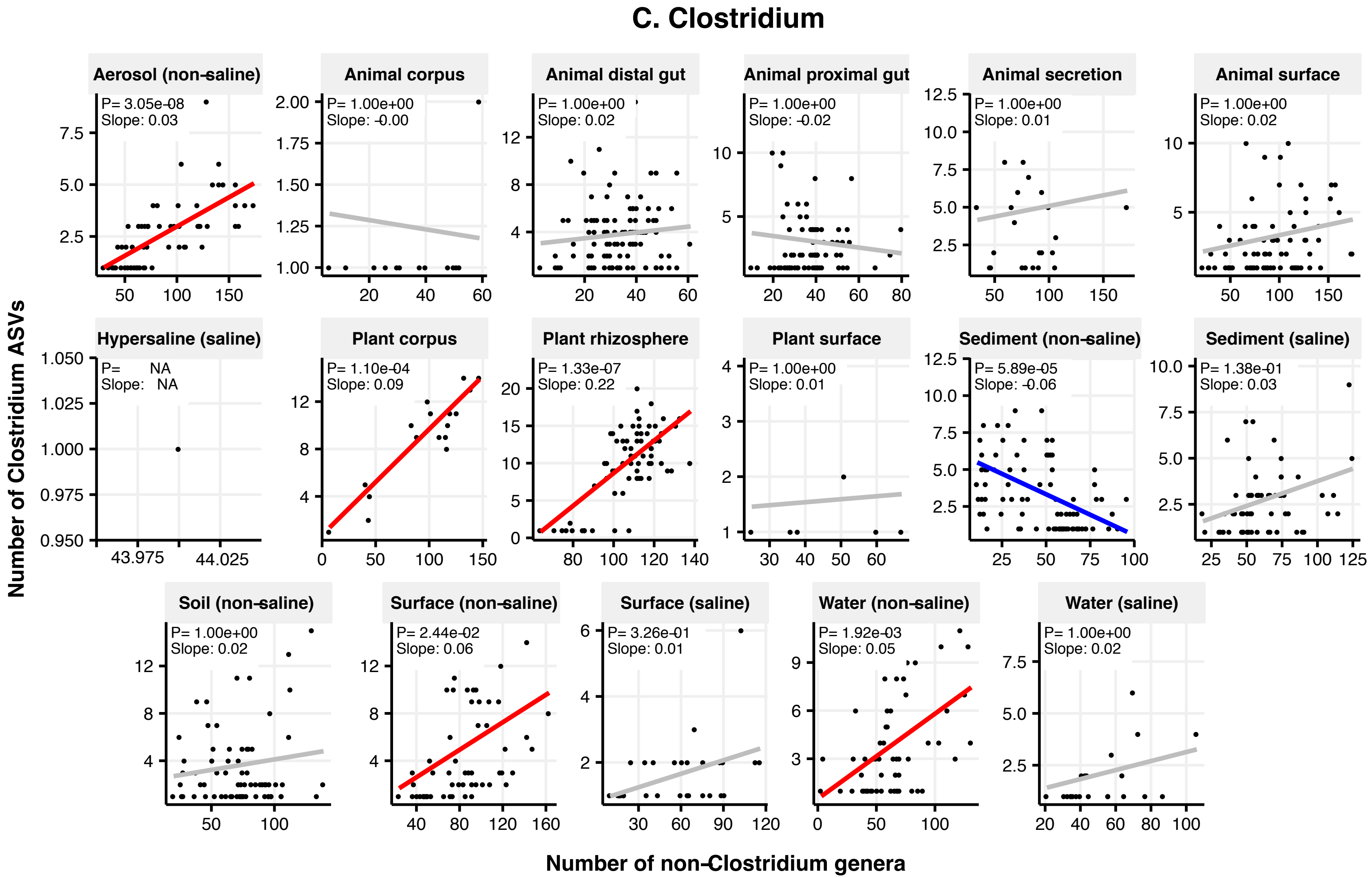

### Figure 2 Supplement 7

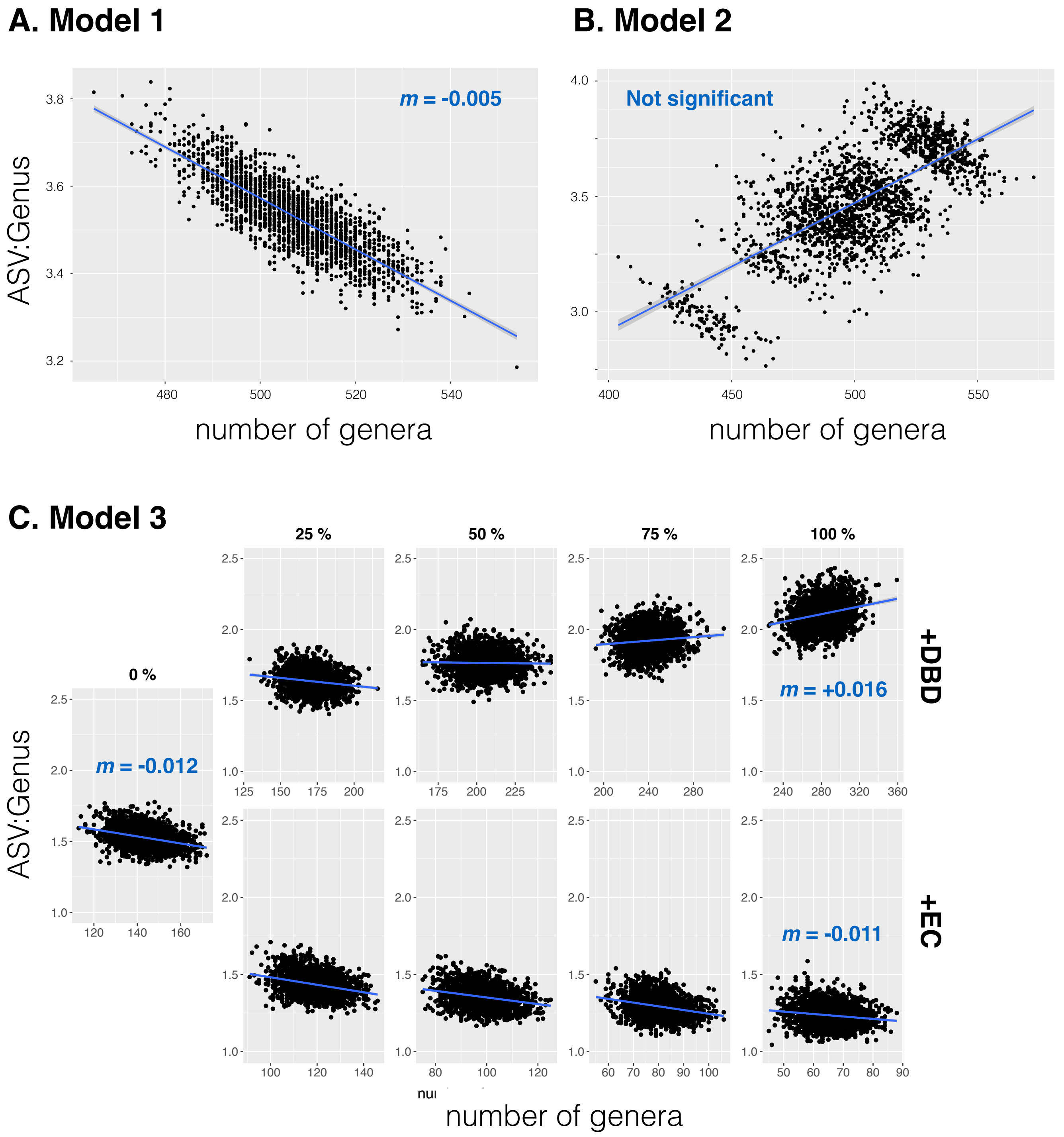

### Figure 2 Supplement 8

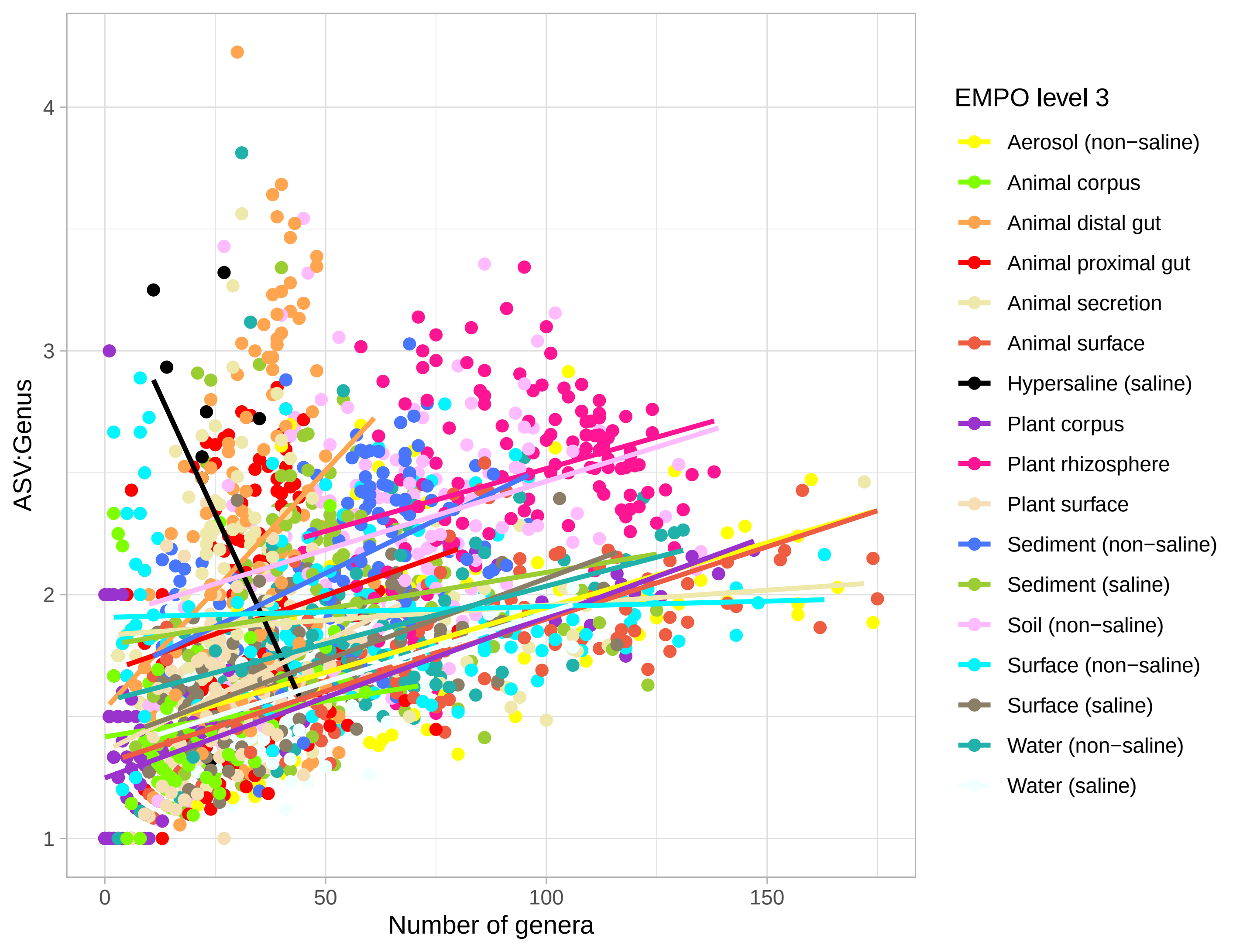

### Figure 2 Supplement 9

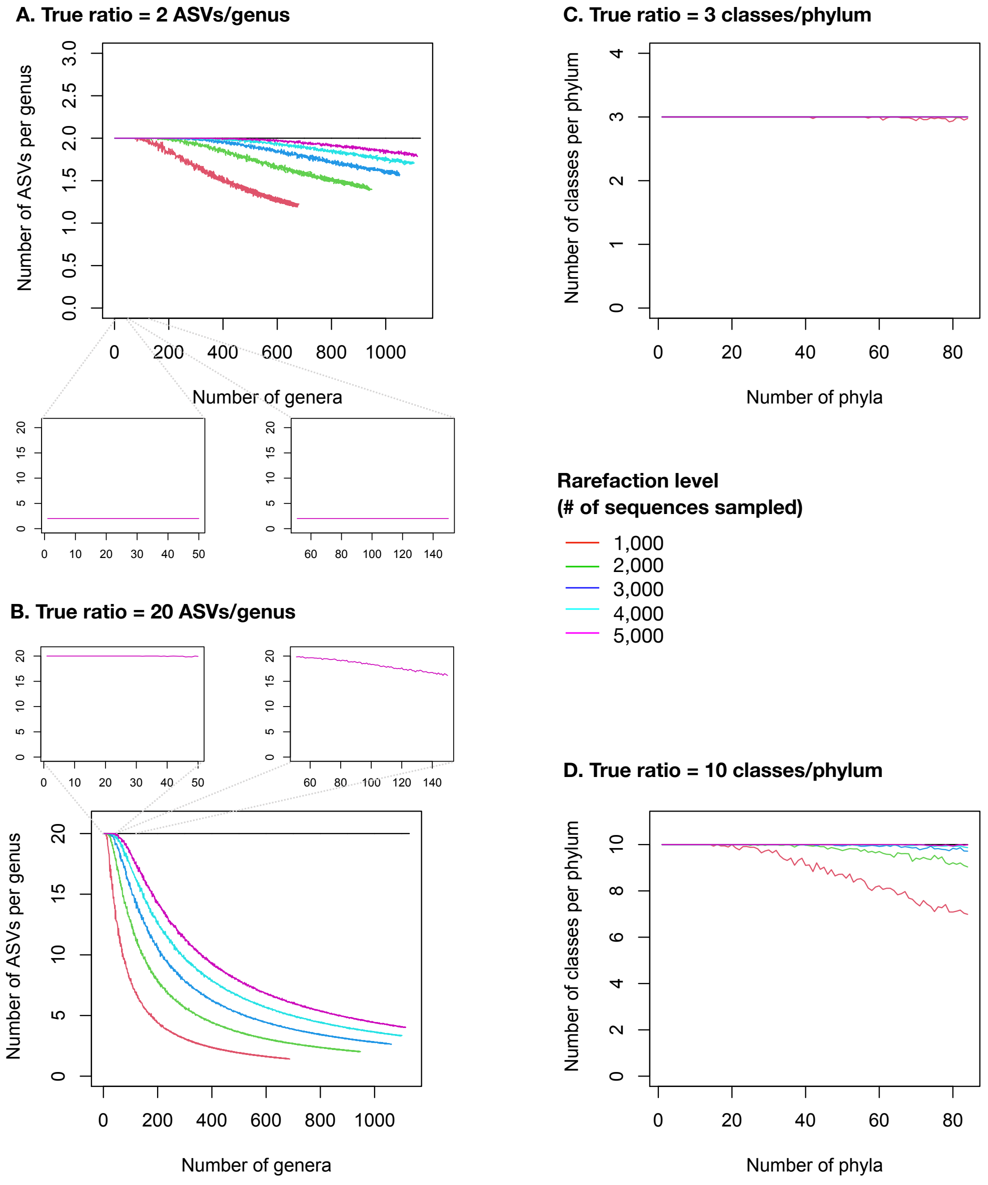

### Figure 2 Supplement 10

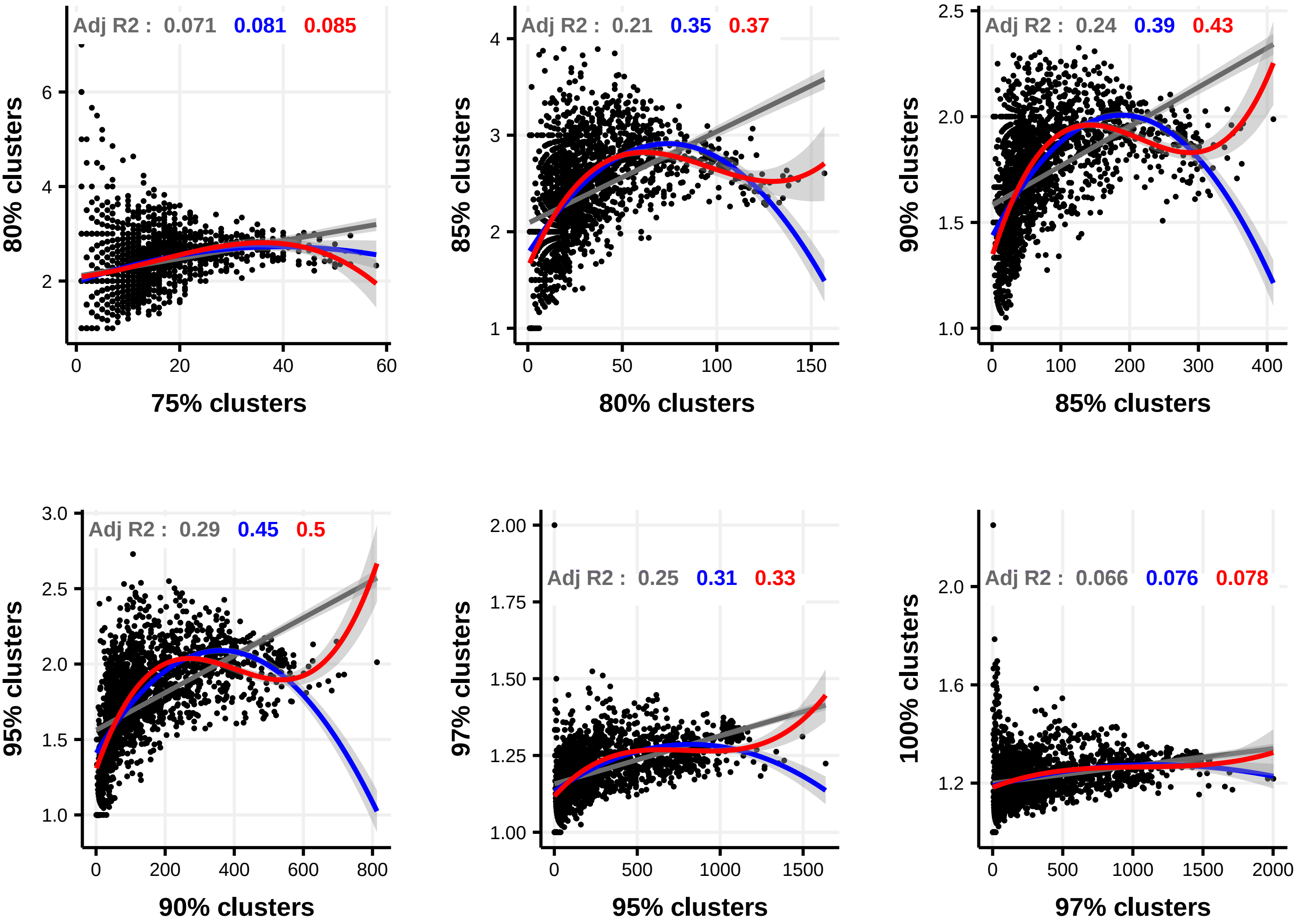

### Figure 2 Supplement 11

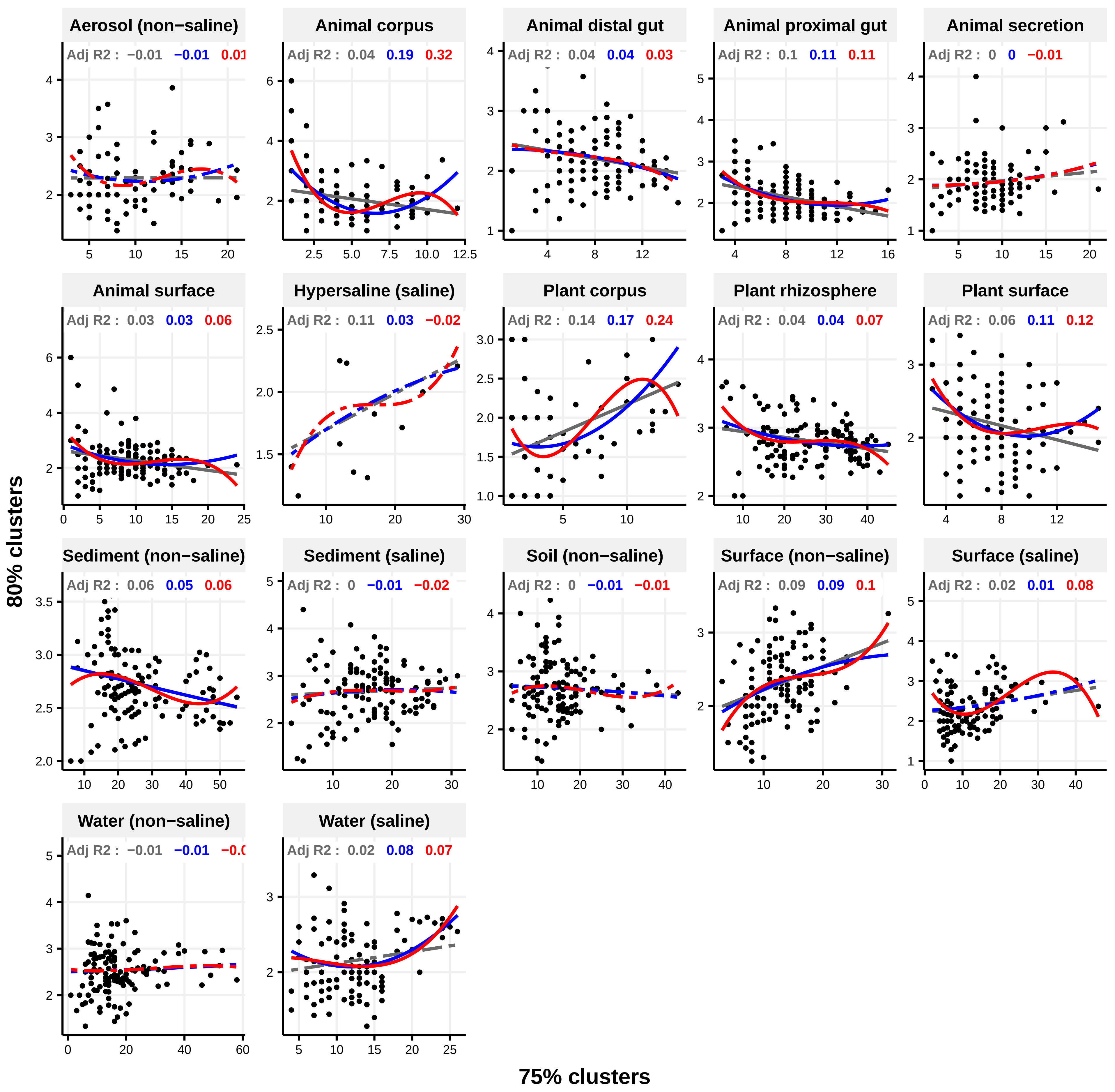

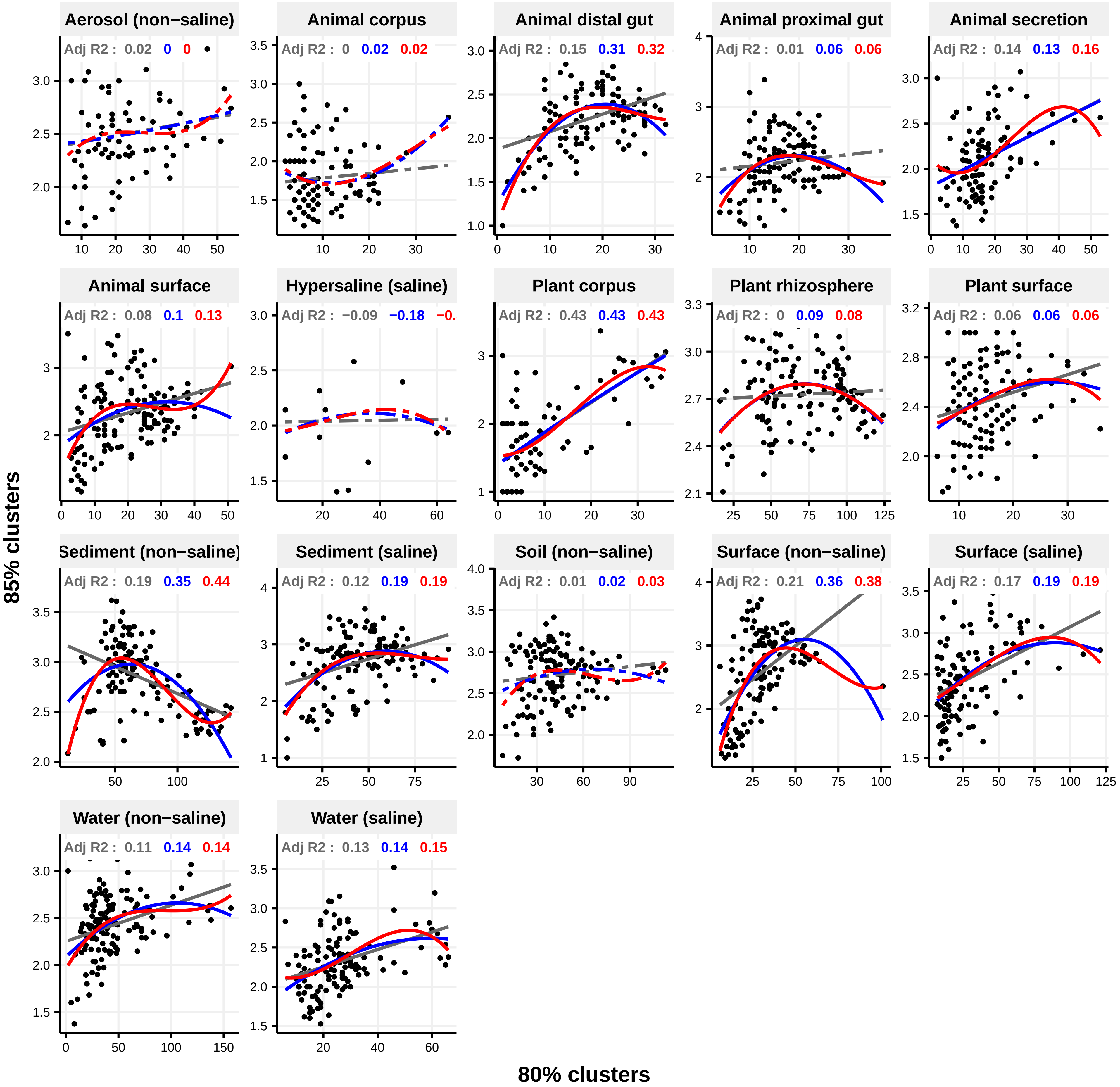

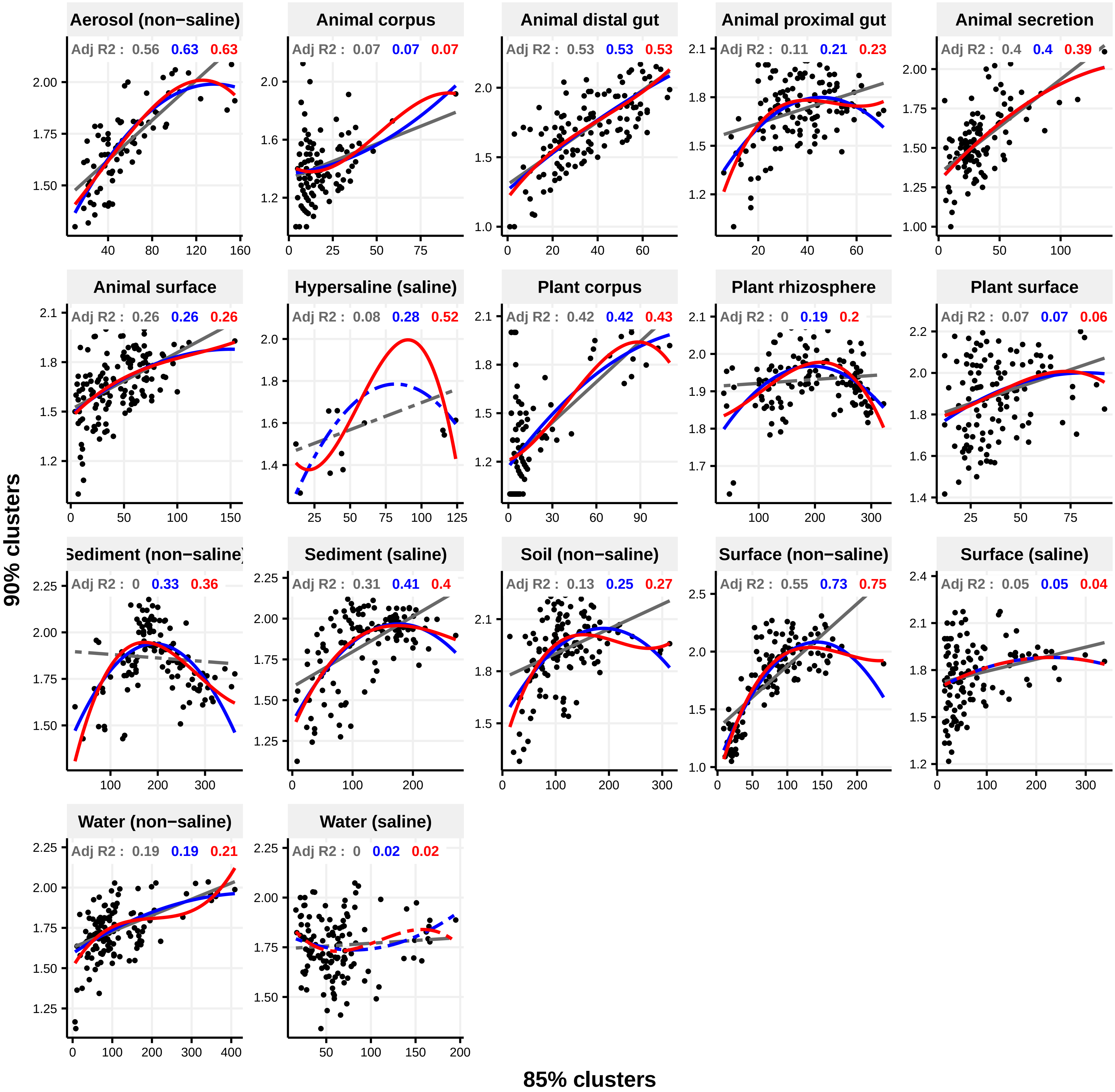

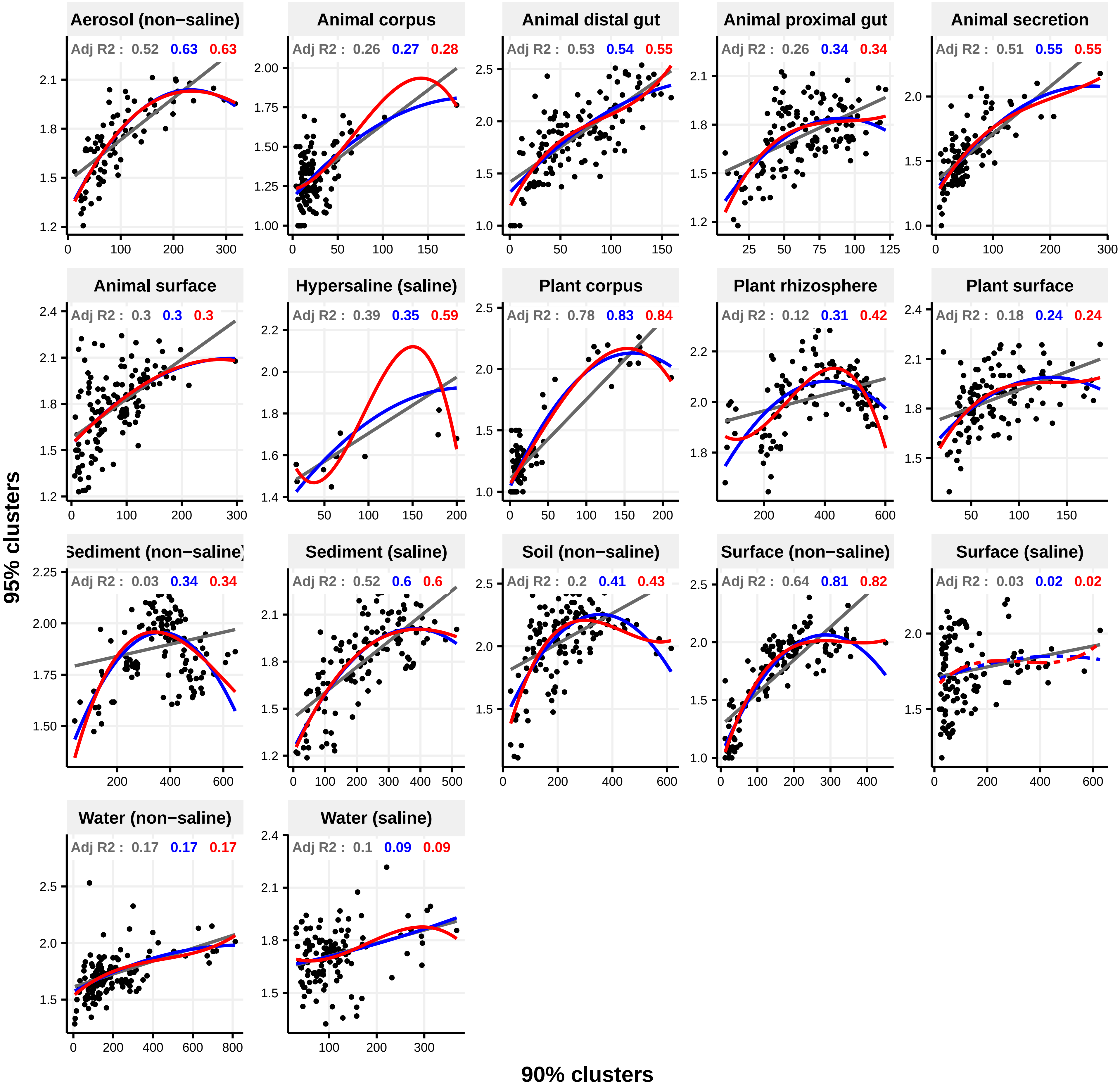

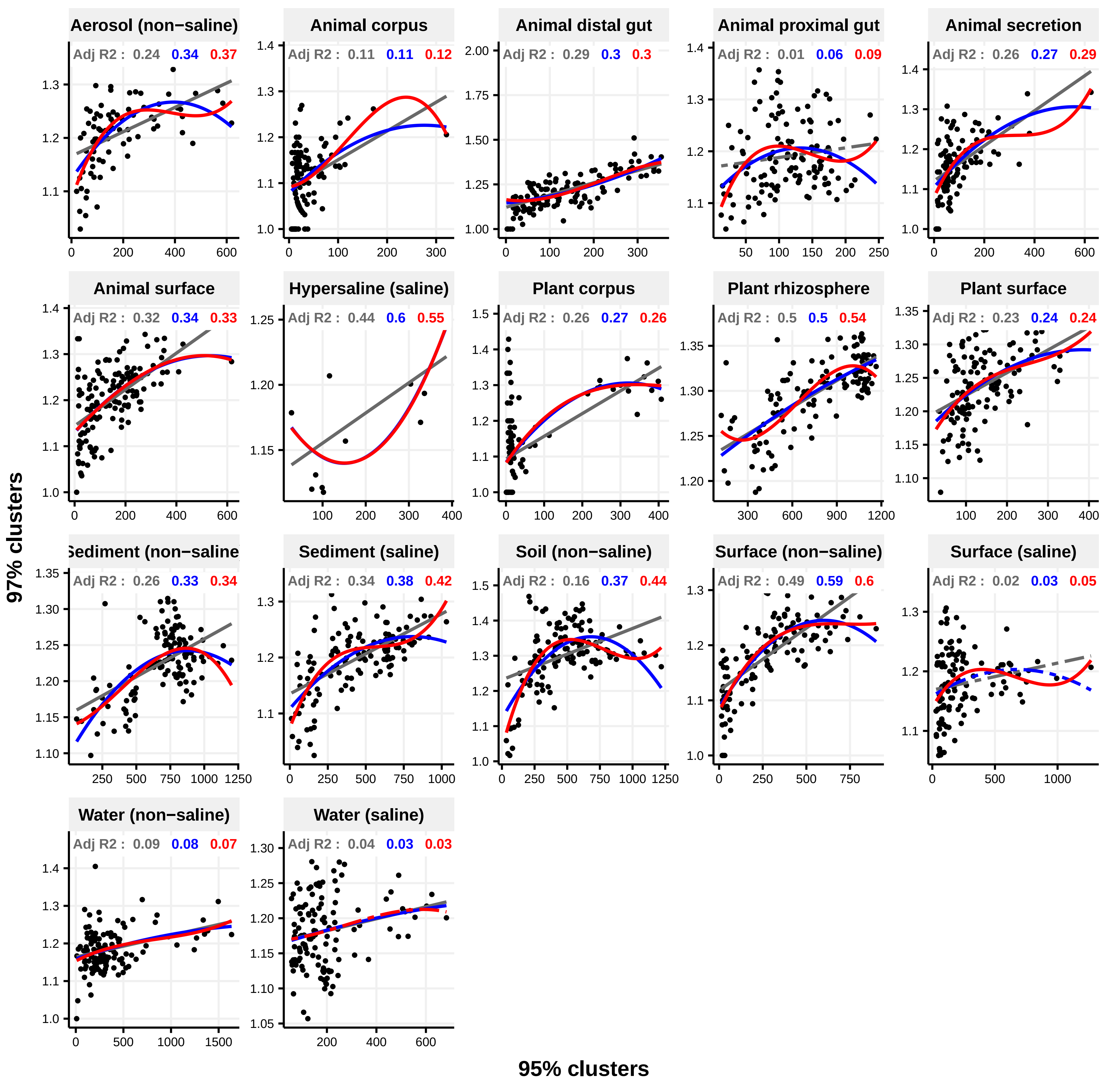

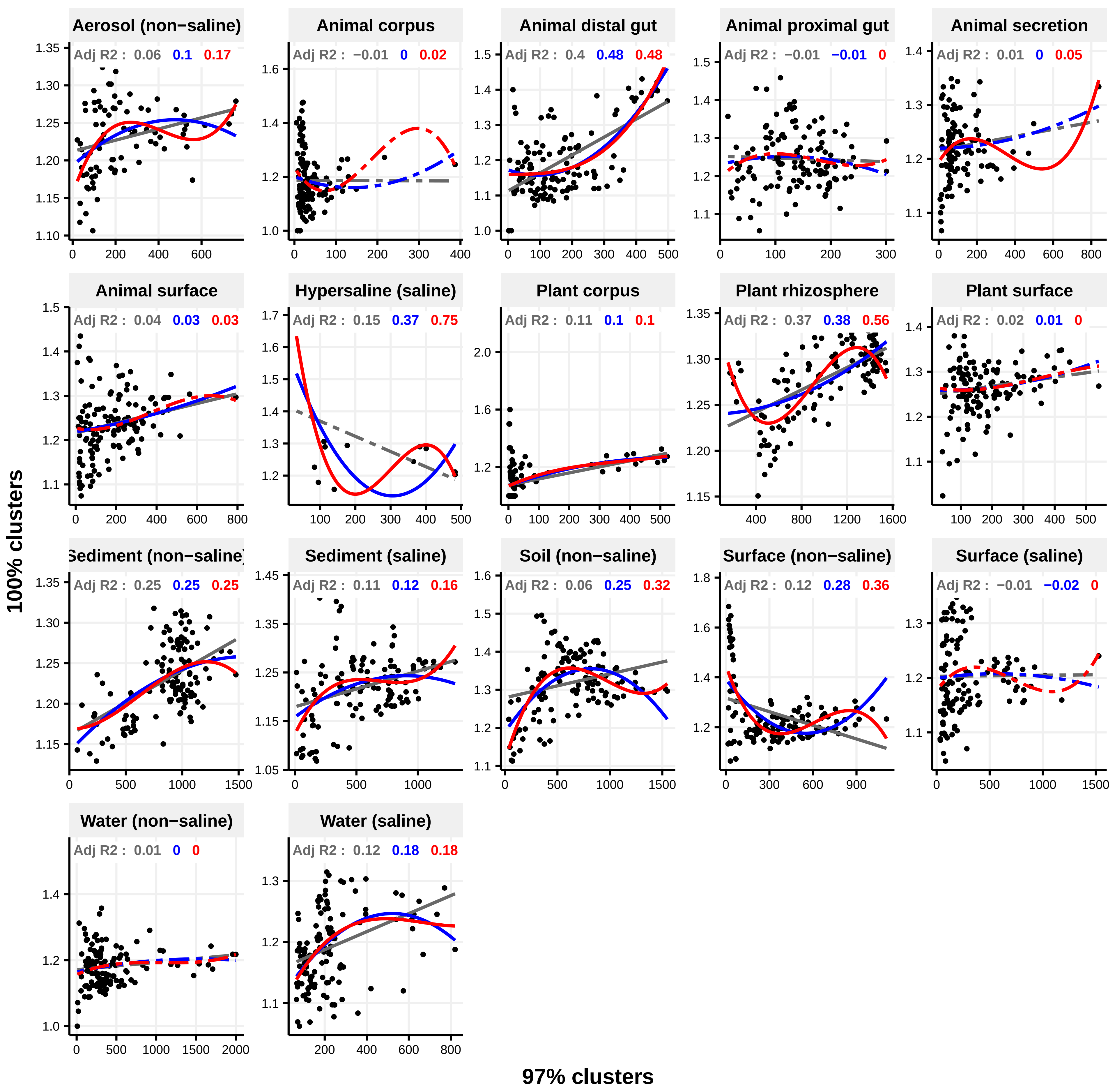

### Figure 4 Supplement 1

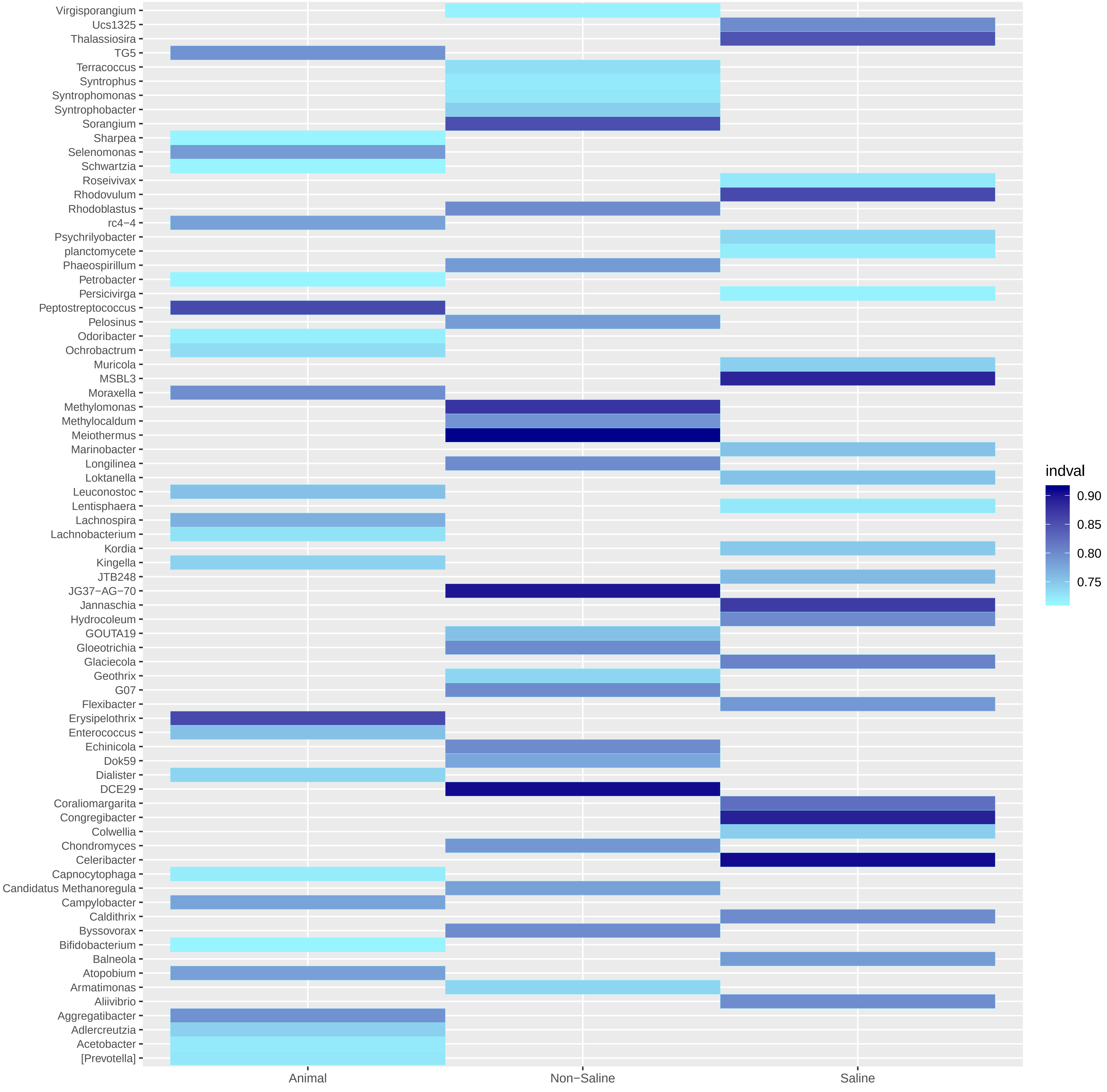
